# ConvexGating infers gating strategies from clusters in single cell cytometry data

**DOI:** 10.1101/2024.11.11.623019

**Authors:** Vincent D. Friedrich, Karola Mai, Thomas P. Hofer, Elfriede Nößner, Lorenzo Bonaguro, Celia L. Hartmann, Aleksej Frolov, Caterina Carraro, Doaa Hamada, Mehrnoush Hadaddzadeh-Shakiba, Markus Scholz, Fabian J. Theis, Marc D. Beyer, Joachim L. Schultze, Maren Büttner

**Affiliations:** University of Leipzig, Institute for Medical Informatics, Statistics, and Epidemiology, Leipzig, Germany; Center for Scalable Data Analytics and Artificial Intelligence (ScaDS.AI), Leipzig, Germany; Systems Medicine, Deutsches Zentrum für Neurodegenerative Erkrankungen (DZNE), Bonn, Germany; Modular High Performance Computing and Artificial Intelligence, German Center for Neurodegenerative Diseases (DZNE), Bonn, Germany; Research Group Tissue Control of Immunocytes, Helmholtz Center Munich, Munich, Germany; Life and Medical Sciences (LIMES) Institute, University of Bonn, Bonn, Germany; PRECISE Platform for Single Cell Genomics and Epigenomics, DZNE and University of Bonn and West German Genome Center (WGGC), Bonn, Germany; Immunogenomics & Neurodegeneration, Deutsches Zentrum für Neurodegenerative Erkrankungen (DZNE), Bonn, Germany; Molecular Immunology in Neurodegeneration, German Center for Neurodegenerative Diseases and the University of Bonn, Germany; Medical Microbiology and Immunology Department, Faculty of Medicine, Mansoura University, Egypt; University of Leipzig, Faculty of Mathematics and Computer Science, Leipzig, Germany; Institute of Computational Biology, Helmholtz Center Munich, Germany; Department of Mathematics, Technical University of Munich, Germany; TUM School of Life Sciences Weihenstephan, Technical University of Munich, Germany

## Abstract

Manual expert gating remains common practice for the definition of specific cell populations in the analysis of flow cytometry data. The increasing number of measured parameters per individual cell and high inter-rater variability makes manual gating inconsistent in many scenarios such as multi-center studies. Here, we propose ConvexGating, an AI tool that automatically learns gating strategies in an unbiased, fully data-driven, yet interpretable manner. ConvexGating scales efficiently with increasing parameter space, creating proficient strategies with low-contamination in the extracted population for previously known and so far unknown or ill-defined cell populations. The inferred strategies are independent of parent populations, for instance, plasmacytoid dendritic cells (pDCs) can be fully identified as CD45RA- CD123+. In addition to flow cytometry data, ConvexGating derives gating strategies for cyTOF (Cytometry by Time of Flight) and CITEseq (Cellular Indexing of Transcriptomes and Epitopes by Sequencing) data and supports optimal design of marker panels for cell sorting.

## Introduction

Flow cytometry is a widely used method for analyzing biological properties of single cells as well as sorting of cell populations for functional testing. Recent machine- and fluorochrome-wise technological advances enabled the measurement of up to 30 cellular markers in parallel. This leads to the acquisition of more and more extensive cellular signatures in flow cytometry experiments. In current mass cytometry (cyTOF) applications commonly 40 and more markers are used^1,2^ with a usually lower cell throughput than flow cytometry^2^. In sequencing-based readouts for antibody binding to cellular protein markers (e.g. CITEseq), antibody panels utilizing even up to 200 different markers have been reported^3,4^. Consequently, the use of machine learning based analysis methods is more common for high-dimensional mass cytometry and CITE-seq data^5^.

For the analysis of flow cytometry data is manual gating, a stepwise procedure of selecting cell subsets based on combinations of expression or intensities of characteristic cellular markers, still common and best practice. In particular, one inspects the distribution of two selected markers and draws a gate around the population of interest in the corresponding two-dimensional space. Subpopulations within the gate are further distinguished by iteratively examining other marker pairs. This approach becomes increasingly laborious when up to 30 markers are analyzed. Furthermore, manual gating strategies are subject to variations across operators^6–8^ and over time for a single operator^8–10^. For instance, the separation of monocytes into classical, non-classical and intermediate subpopulations showed considerable inter-rater variability^7^.

In contrast, clustering methods like PhenoGraph^11^ or Leiden clustering^12^, amongst others^13^, are most common for analyzing mass cytometry data, where panels encompass up to 40 markers. These unbiased approaches consider all markers simultaneously, which results in a more flexible separation of cells based on multiple features at once. In contrast to manual gating, clustering methods label all cells in a data set. To facilitate high-dimensional data analysis for flow or mass cytometry data, we previously developed the package *pytometry*^14^, which extends the popular single-cell framework scanpy^15^ and builds upon the annotated dataframe (anndata)^16^ data structure and related Python packages from the scverse^17^. This integrative approach also paves the way to popularize machine learning applications for flow and mass cytometry data. Such more comprehensive analysis can better identify specific subpopulations and discover novel cell types that might be missed using traditional methods. For instance, Georg et al^18^ analyzed cyTOF data from COVID-19 patients and identified disease specific CD4+ and CD8+ T cells that expressed CD16 via clustering. The abundance of these T cells was associated with severe COVID-19 outcome. As a second example, basophils lose the basophil-defining marker CD123 upon activation^19^, which renders the recovery of basophils in the clinical basophil activation test more difficult. Accordingly, the discovery of novel cell types typically benefits from clustering to identify cell subtypes and states and further improves by contextualization with healthy reference data. In addition, clustering offers more flexibility to jointly analyze multiple samples at once, thereby facilitating multicenter studies.

Clustering also provides a more comprehensive view on the variability of innate immune cells. In particular, it was demonstrated that cell typing and lineage tracing benefit from high-dimensional data spaces and unbiased clustering outperformed manual gating based analysis^20–22^. For example, the identification of plasmacytoid DCs (pDCs) can be compromised by precursors of conventional DCs (pre-cDCs) when only few markers are used^20,23,24^. As a result, a cell population sorted by flow cytometry could be affected by a potentially unknown level of cross-contamination. Such a population of cells may respond heterogeneously to a stimulus, either due to the inherent heterogeneity of the population or due to being a mixture of different cell types. Identifying the underlying cause for the differences in the response is then challenging.

Still, manual gating remains gold standard, also due to the lack of complementary control experiments to determine true cellular identity. Recent advances on multimodal experiments like CITE-seq^3^, Abseq^4^ or REAP-seq^25^ combine surface antibody labeling and gene expression profiling at single-cell resolution. Such multimodal data provide independent information on cellular identity required to compare manual gating with clustering and other machine learning-based classification tools.

Using clustering to identify cell population has several advantages, however, transferring the cell identity back into a sorting strategy for further experiments requires reverse engineered gating strategies. An optimal gating strategy aims to recover the cells of interest (recall) with high purity (precision) in as few steps as possible. While clustering can inform the construction of a gating strategy, it largely depends on manual finetuning. This process can be significantly enhanced by machine-learning (ML) approaches. A more recent approach called Hypergate^26^ identifies populations via shrinkage and expansion of rectangles in the full marker space. Ji et al.^27^ present an approach for learning rectangular gate boundaries through gradient descent. However, certain subpopulations might require more flexible gate geometries than rectangles. Especially to avoid contamination with undesired non-target cells for small cell subpopulations, an arbitrary flexible gate shape in 2D marker space is beneficial. Enforcing a convex gate shape ensures a continuous gated region, thus sacrificing a certain level of performance for interpretability and reproducibility which can therefore be considered a form of regularization. To this end, we developed ConvexGating to derive full gating strategies for any cell population of interest in an autonomous fashion independent of manual expert gating. The suggested gating strategies aim at retrieving the cell populations of interest as much as possible while concomitantly avoiding contamination with non-target cells.

ConvexGating proposes gating strategies consistent with manual gating and outperforms existing tools, especially for fine-grained cell types. ConvexGating works equally well with the full range of panel sizes from 10 markers in FACS panels up to 200 markers in CITE-seq panels. For previously unknown populations in COVID-19, such as CD16+ T cells, we illustrate for ConvexGating to reliably infer gating strategies across multiple modalities and patient cohorts.

## Results

### ConvexGating ML workflow mimics manual gating

Manual gating is an iterative process, where users select a set of markers to separate a population of interest from the remaining cells based on prior knowledge. In our approach, ConvexGating, we mimic manual gating in an automatic, fully data-driven way. Our approach is based on the assumption that every cell population in a flow or mass cytometry dataset can be identified by an appropriate clustering^11,12,28^ (**Fig. 1 A**).

**Figure 1:**
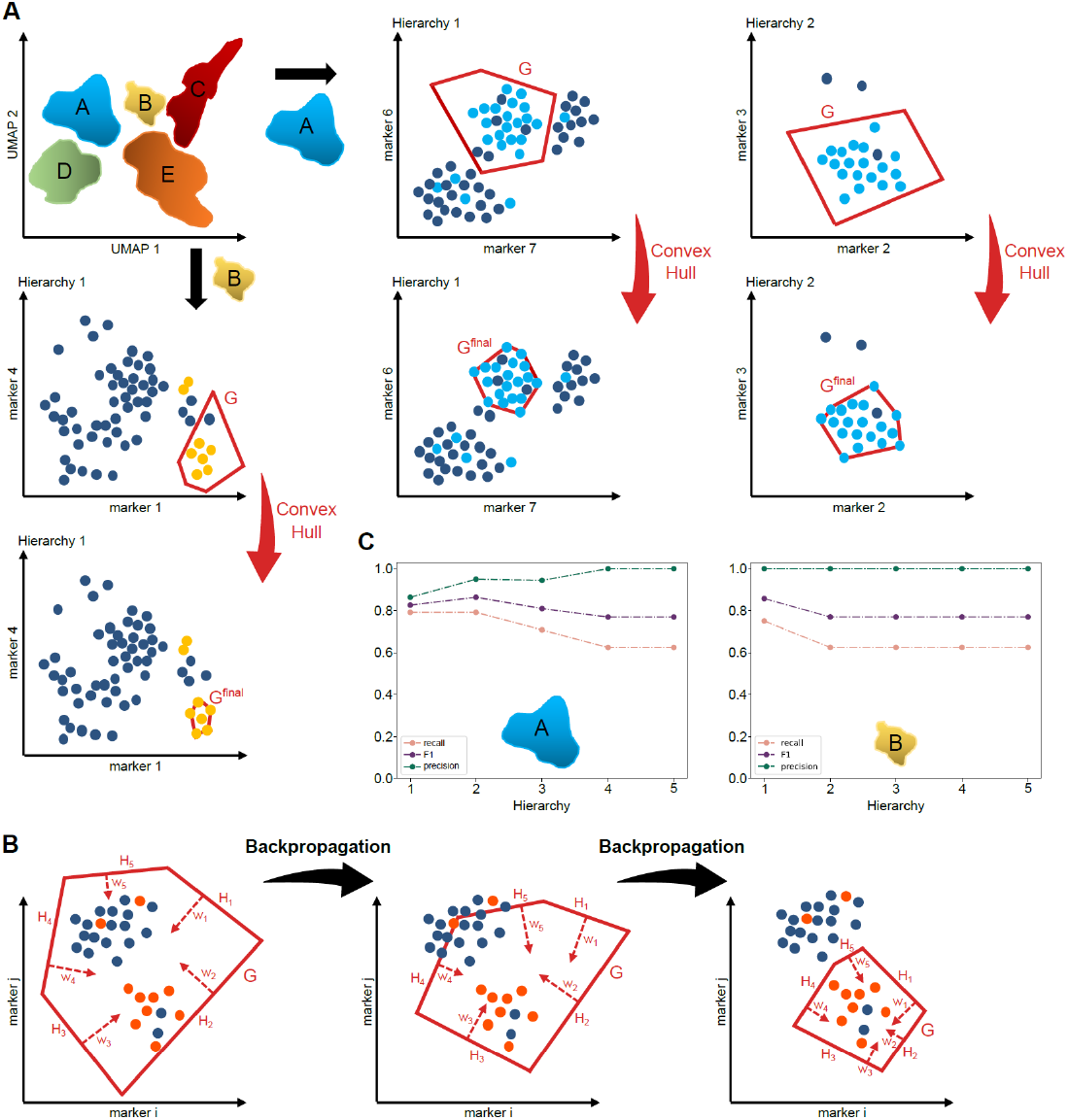
ConvexGating schematic workflow. **(A)** ConvexGating requires labeled input. Colors in 2D UMAP space reflect assigned cell labels (top left). Per gating hierarchy, ConvexGating identifies the two features that exhibit the largest difference in distribution between targets and non-targets. After mapping into the corresponding 2D marker space ConvexGating learns the optimal location of an intermediate gate G in 2D marker space by trading-off recall and precision via an explicitly designed loss function. Shrinkage of gate G via the convex hull of targets inside G increases precision and yields final gate *G*^*final*^. Exemplary gating strategy for cell population A with two hierarchies (top right) and for cell population B with one hierarchy (down left). **(B)** Learning optimal gate location in 2D marker space to separate target cells (orange dots) from non-target cells (blue dots) via stochastic gradient descent (SGD). Gate G (red) is defined as the intersection of several 2D halfspaces that are each specified by a 2D normal vector and a 1D bias parameter. Gate G gets randomly initialized with some of the normal vectors fixed based on the principal components of the target population (left). The location of gate G is iteratively adapted by back-propagating the error of a version of binary cross-entropy loss complemented by two problem-specific weighted regularization terms. The normal vectors and bias parameters that specify gate G are updated accordingly (middle). The SGD-like gate updates continue until a termination criterion is satisfied (right). **(C)** ConvexGating is evaluated by recovery of target cells (recall), by contamination with other cell types (precision), and by the weighted mean of precision and recall (F1 score). Maximum F1 score determines the number of hierarchies in the final gating strategy.

We define the population of interest as the target population and refer to all other cells as the non-target population. In the first step, we calculate distribution parameters along each marker to select the most promising marker combination in terms of separability of target and non-target population in the current gating hierarchy (see **methods**). Next, we represent the gate surrounding the target population as a convex shape bounded by a set of hyperplanes. One subset of hyperplanes is randomly initialized while the normals of the remaining hyperplanes are fixed based on the principal components derived from the data matrix of the target population. During the optimization step, the hyperplanes form an intermediate gate by minimizing a modified version of binary cross-entropy loss complemented by two problem-specific weighted regularization terms using stochastic gradient descent (**Fig. 1 B**). This loss incentivizes balancing recovery of target cells (recall) and purity of the gated population (precision). In the final step, we tighten the intermediate gate(s) to exclude as many non-target cells as possible. This is achieved by applying the convex hull to the encircled target population within the intermediate gate(s).

We then repeat the process of obtaining an optimal marker combination and subsequent gate optimization until we obtain the cleanest possible population (high precision) and retain as many cells as possible (high recall) (**Fig. 1 C**). ConvexGating prioritizes precision over recall, aiming for purity in the extracted population. We use F1 scores as measure for the balance between precision and recall and monitor these metrics for all gating hierarchies. The final gating strategy retains hierarchies up to the level where the highest F1 score occurs.

### ConvexGating infers gating strategies similar to manual gating

In our first showcase, we demonstrate that cell type annotations achieved through clustering and manual gating are highly consistent, and inferred gating strategies successfully identify all target populations with high precision. In human blood, there are three dendritic cell (DC) populations classified as plasmacytoid DCs, CD141+ myeloid DCs, also termed cDC1, and CD1c+ myeloid DCs, also termed cDC2. Classification is not trivial as there is no lineage marker. Consequently, markers are shared with other blood leukocytes and marker expression depends on the activation state of the cells. Focusing on the CD1c+ DC2, there is a CD14+ and CD14- population^29^ and a differential expression of CD5 has been described^30^. Hence, there are at least two subsets of CD1c+ DC2 cells, which are CD1c+ CD5- CD14- cells, CD1c+ CD5+ CD14- cells, and CD1c+ CD5- CD14- cells. Current data suggest that the CD14- cells are bona fide DCs, while the CD14+ cells represent a unique population with ontogenetic links to monocytes^31^. Therefore, the correct dissection of the CD14+ and CD14- populations is of great importance. Care must be taken, as CD14 is expressed by all three monocyte subsets and the CD1c+ CD5- CD14+ DC2s tend to overlap with the classical monocytes. Also, CD14 is expressed at low levels by neutrophils, and CD1c+ is also expressed on B lymphocytes. To develop a robust identification strategy for the cDC2 subsets, we assembled a 6-marker panel including CD1c, CD5, CD14, CD16, CD19/20, and HLA-DR. We will refer to this panel as the DC panel in the following. In whole blood stainings from three healthy donors, we used both manual gating (**Fig. 2 A** and **Supplementary Fig. 1**) and Leiden clustering with our previously developed Python package *pytometry*^14^ (**Fig. 2 B** and **Supplementary Fig. 2-5**) to identify B cells (CD19/20+), monocytes (HLA-DR+, CD14+ or CD16+) and type 2 dendritic cells (cDC2s) (HLA-DR+, CD1c+). Notably, T cells remained unannotated as the panel did not include CD3 to stain T cells. We further subtyped classified monocytes into classical monocytes (CD14+, CD16-), non-classical monocytes (CD14low, CD16+) and intermediate monocytes (CD14+, CD16+) through clustering (**Supplementary Fig. 3**). For the identification of cDC2 subsets, we distinguished CD1c+ DC2s based on CD5 and CD14 markers as described above^31^ (**Supplementary Fig. 4**). For both monocytes and cDC2 subtyping, we observed less overlap between manual gating and clustering, while the overall overlap with the manual gating annotation is high (see **methods**) (**Fig. 2 C**). In case of monocyte subtypes, we attribute the lower concordance to the continuous decrease of CD14 and the increase of CD16, which make the distinction of the subpopulations difficult. For cDC2s subpopulations, we work with relatively few cells resulting in less concordance between manual gating and clustering.

**Figure 2:**
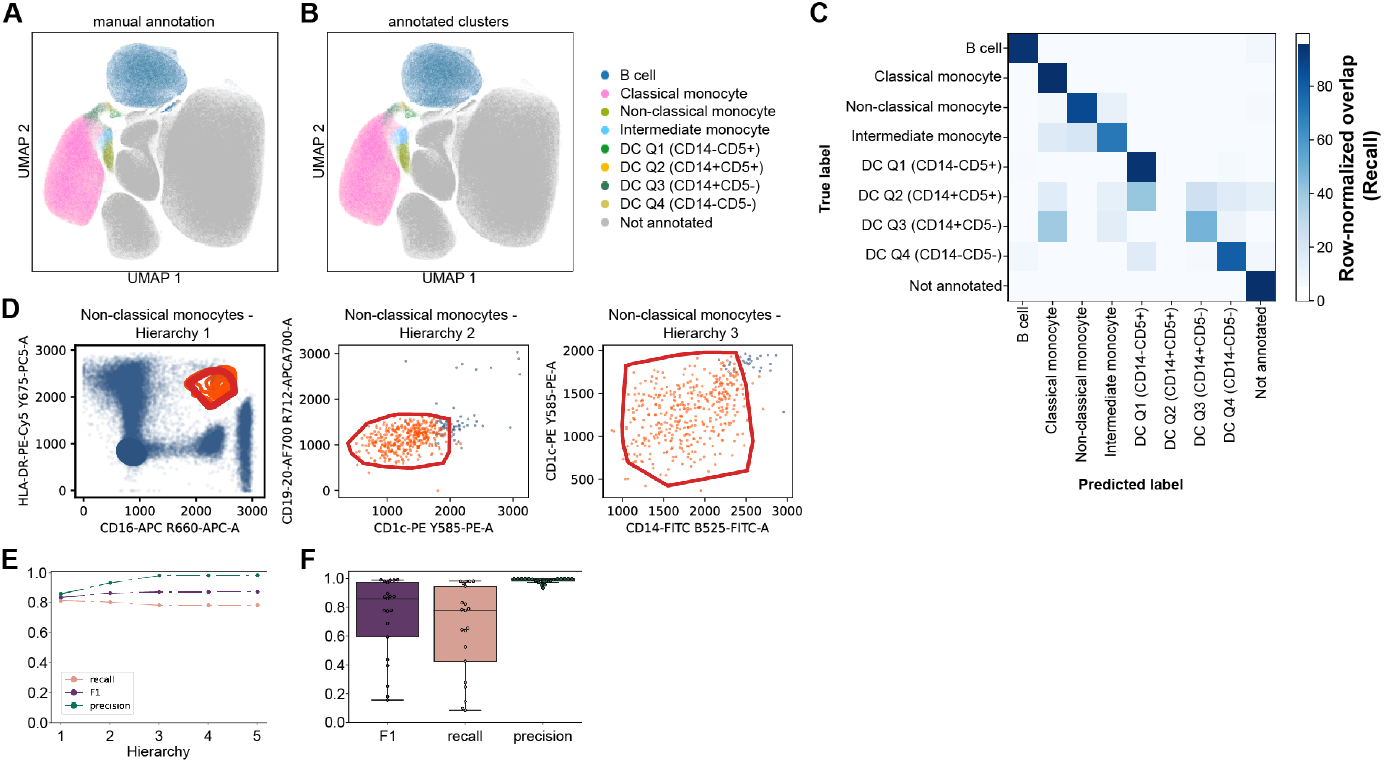
ConvexGating infers gating strategy for annotated clusters on a monocyte panel. **(A)** UMAP displaying expert cell type annotation via manual gating. **(B)** UMAP displaying cell type annotation via clustering. **(C)** Row-normalized overlap (recall) of clustering with expert labels. **(D)** Gating strategy for non-classical monocytes inferred by ConvexGating. Target cells (orange) and non-target cells (blue) are represented via scatter plots and/or contour plots per target and non-target population. **(E)** Precision, recall and F1 score for each hierarchy in C). No improvement in F1 score in higher hierarchies observed. **(F)** Boxplot of precision, recall and F1 scores for all annotated cell populations.

We then applied ConvexGating to all cell types identified by clustering and examined the proposed gating strategies (**Fig. 2 D** and **Supplementary Fig. 6 A-I**). Here we showcase the ConvexGating results for the case of non-classical monocytes (**Fig. 2 D**). The optimal gating strategy consists of three hierarchies and largely reflects the manual gating strategy (CD19/20-, HLA-DR+, CD16+, CD14-). In every hierarchy, we observe an increase in both F1 score and precision with final values of 0.98 for precision, 0.79 for recall and 0.87 for F1 score for this population (**Fig. 2 E**). As optimized for highest precision, over all populations, our computational gating strategy has very high precision scores and for most cell types also high F1 scores (**Fig. 2 F** and **Supplementary Fig. 6 D**). Taken together, ConvexGating inferred precise and plausible gating strategies for abundant and rare cell types.

### Considerable variability exists across operators in a manual gating task

Next, we investigated the robustness of manually acquired gates on a larger flow cytometry panel with 27 markers using 5 frozen PBMC samples of healthy donors^32^. Specifically, we devised the task to gate for major cell types, such as T cells, B cells, NK cells and monocytes, as well as a finer resolution of T cell, monocyte and DC subpopulations (see **methods**) (**Fig. 3** and **Supplementary Fig. 7 A**). We obtained a total of 10 submissions from 7 operators with different levels of expertise. As additional submission, we pre-processed the same datasets computationally with debris removal (QC) and identification of cell populations with *pytometry* using Leiden clustering. We defined the gating result of the original study^32^ as the Gold standard for our comparison. To evaluate the concordance among operators, we first analyzed which cells are classified as debris (including doublets) and valid cells in the QC using FSC and SSC, and then measured the overlap with the Gold standard using the Jaccard score (**Fig. 3 A**, see **methods**). Seven submissions marked almost identical cells as valid cells, while two submissions were more restrictive, resulting in lower Jaccard scores. The computational QC recovered slightly less cells compared to the Gold standard (**Fig. 3 A** and **Supplementary Fig. 7 B**), but overlaps largely with gating-based QC. In the Gold standard around 89% of events passed QC as valid cells. Over all submissions, 9.48% of events were consistently labeled as debris (**Fig. 3 B**) and we did not observe a QC bias over samples (**Supplementary Fig. 7 C**).

**Figure 3:**
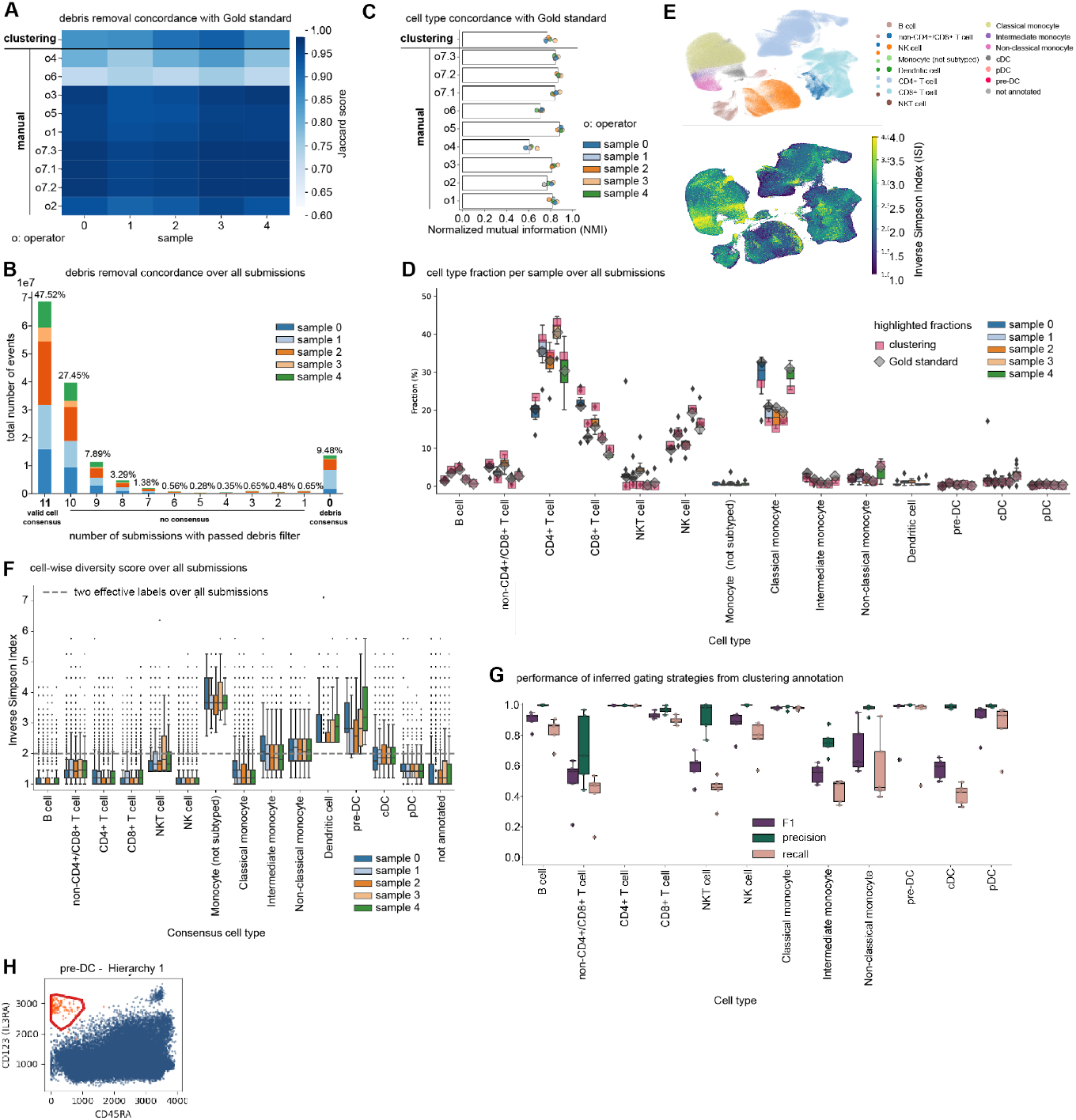
Assessing the variability of human expert labeled data on a high-dimensional marker panel. **(A)** Heatmap depicting debris removal across all submissions compared to Gold standard. Coloring indicates Jaccard score over all cells per sample **(B)** Stacked barplot on the number of cells labeled as valid aggregated over all submissions, colored by sample. **(C)** Barplot of overlap of cell type labels compared to Gold standard measured by normalized mutual information (NMI). Dots colored by sample indicate NMI per sample. **(D)** Boxplot of cell type composition over all submissions (n=11, boxes represent 25th percentile, median and 75th percentile, respectively, whiskers 1.5 * interquartile range and black diamonds represent outliers), colored by sample. Pink squares indicate cell type annotation by clustering, gray diamonds indicate Gold standard annotation. **(E)** UMAPs displaying the consensus cell type labels over all submissions and Gold standard (upper panel) and the number of effective labels per cell over all human expert labels measured by Inverse Simpson Index (ISI) (lower panel). **(F)** Boxplot of number of effective labels per cell over all submissions (n=11, boxes represent 25th percentile, median and 75th percentile, respectively, whiskers show 1.5 * interquartile range and black diamonds represent outliers), colored by sample, grouped by the consensus label over all submissions and the Gold standard. Grey dashed horizontal line indicates the effective number of 2 cell type labels. **(G)** Boxplots summarizing precision, recall and F1 scores per sample (n=5, boxes represent 25th percentile, median and 75th percentile, respectively, whiskers show 1.5 * interquartile range and black diamonds represent outliers) obtained by running ConvexGating on all samples for cell populations annotated by clustering. **(H)** One-step gating strategy for pre-DCs inferred by ConvexGating.

Next, we quantified the robustness of cell type labels with several metrics. Firstly, we evaluated their overlap with the Gold standard using normalized mutual information (NMI) (**Fig. 3 C** and **Supplementary Fig. 7 D** see **methods**). The overlap of cell types and their compositions is largely consistent across all submissions (**Fig. 3 C**), and the clustering results are within the range of observed variability of the results. Secondly, we computed the variability of cell type compositions (**Fig. 3 D** and **Supplementary Fig. 7 E**). Specifically, we highlighted the compositions obtained via clustering and by the Gold standard as gray squares and diamonds, respectively, superimposed on the boxplot of cell type compositions. Overall, the deviation of clustering to the Gold standard lies within the variation observed over all submissions (manual gating) and samples, except for assigning slightly more cells to CD4 and CD8+ T cells, while slightly less cells are labelled as classical monocytes. The discrepancy can be explained by the hierarchical approach of manual gating; the T cell gate is usually drawn first while monocytes are identified thereafter. Monocytes ending up as contamination of the T cell gate cannot be re- assigned to the monocyte populations. This highlights the strength of high-dimensional clustering for cell type analysis as it reduces the bias of the hierarchical gating approach. Thirdly, to derive reliable gating strategies, we must ensure that the population of interest has a high purity and actually contains the cells of interest. However, some populations are inherently difficult to identify using gating. Therefore, we determined the number of effective cell type labels per cell using inverse simpson index (ISI) (**Fig. 3 E-F** and **Supplementary Fig.s 7 F-G** see **methods**), where an effective number of 2 or more indicates discordance of the assigned labels across the expert gating strategies. In this assessment, we used the most often assigned label as a consensus label instead of relying on the Gold standard label alone (**Fig. 3 E**). While the overall concordance of cell type labels measured using NMI is high (**Fig. 3 C** and **Supplementary Fig. 7 D**), we observe ISI of 2 and above in labeling DCs and the DC subpopulations cDCs and pre-DCs, as well as intermediate and non-classical monocytes (**Fig. 3 E-F** and **Supplementary Fig.s 7 F-G**). Notably, two submissions did not label any pre-DCs. Some submissions labeled classical monocytes partially as NK cells, which is a common error in manual gating (**Supplementary Fig. 7 H-I**). We conclude that the clustering-based cell type annotation is as accurate as a manual gating-based cell type annotation.

Next, we used the clustering-based cell type annotation as input for ConvexGating to infer gating strategies over all samples and determine the variability of the performance scores across the five samples (**Fig. 3 G** and **Supplementary Fig. 7 J)**. The resulting gating strategies recover the majority of cell types with a precision of 0.95 or higher, except for T cells that are neither CD4+ or CD8+, and intermediate monocytes (**Fig. 3 G**, and **Supplementary Fig.s 7 K-L**). ConvexGating shows a lower performance for some rare cell populations like intermediate monocytes (precision 0.71 - 0.9) and cDCs (precision 0.99 - 1 with F1 0.49 - 0.65). On the other hand, we observe a precision >0.9 for other rare populations like non-classical monocytes and pre-DCs (**Fig. 3 H**) across all samples. At the same time, we observed considerable heterogeneity in the assignment of cell type labels for these populations (**Fig. 3 F**), indicating that their recovery is also challenging for classical manual gating strategies.

### ConvexGating supports optimal design of marker panels

To assess the applicability of ConvexGating to high-dimensional mass cytometry data for optimal gating strategies, we applied it to extract cell populations in eight healthy human bone marrow samples measured by mass cytometry^33^ with a 34-marker panel focused on T cell subtyping (“Oetjen cyTOF panel”, **Supplementary Note 1** and **Supplementary Fig. 8**). With increasing cell type granularity, all metrics decrease slightly while the gating depth increases (**Supplementary Fig. 8 A-C**), indicating the increasing difficulty in capturing T cell subpopulations. While ConvexGating used markers like CD4, CD8 and CD197 most often in the initial hierarchies (see “reduced strategy”), it used a greater variety of markers for later hierarchies (see “full strategy”), leading to a larger marker set than that classically used to identify the respective T cell subpopulations (**Supplementary Fig. 8 D-E**). Moreover, ConvexGating directly infers a gating strategy for a target population, regardless of potential parent populations (**Supplementary Fig. 8 F-G**). For example, the sorting strategy for CD4+ TEMRAs involves five steps with sufficiently high precision after three steps (**Supplementary Fig. 8 G-H**). This underscores the higher flexibility of marker choice with ConvexGating and an increased efficiency especially for sorting rare subpopulations directly regardless of potential parent populations.

### ConvexGating outperforms Hypergate regarding high resolution cell type annotations

We benchmarked ConvexGating, the gating tool Hypergate^26^ and off-the-shelf SVM classifiers on three test scenarios (see **methods** and **Supplementary Note 2**). ConvexGating outperformed Hypergate for finer cell type annotations on the DC panel, the large PBMC panel and the Oetjen cyTOF panel, while Hypergate performed slightly better on F1 scores describing broader cell type annotations on the Oetjen cyTOF panel (**Supplementary Fig. 9**). ConvexGating focused on high precision while Hypergate prioritized high recall. When comparing ConvexGating to SVM classifiers, we found that ConvexGating outperformed the linear SVM classifier for finer cell type annotations of the DC panel while the non-linear RBF SVM classifier obtained higher F1 scores for nearly all cell populations (**Supplementary Fig. 10)** at the cost of interpretability of the gate definitions.

### ConvexGating provides gating strategies for disease-specific, previously unknown cell populations associated with severe COVID-19 outcome

ConvexGating creates efficient strategies for previously unknown, ill-defined or perturbed cell populations (**Fig. 4**). Georg et al.^18^ described CD16+ T cells in several independent cohorts of COVID-19 patients. Specifically, the authors reported CD38hi HLA-DR+ Ki-67+ subtypes (clusters 7 and 25) and CD3+ CD16+ highly activated NK-like cells (clusters 8 and 26) in both the CD4+ and CD8+ T cell compartment using cyTOF (**Fig. 4 A**). They confirmed the existence of these populations via manually derived gating strategies. We applied ConvexGating on these populations to identify alternative gating strategies re-identifying these previously unknown and rare populations (**Fig.s 4 B - E**). For the identification of cluster 7, ConvexGating selected CD38 and ICOS in the first hierarchy, and separated further with CD8- and CD28 high (**Fig. 4 B**). Interestingly, cluster 8 was previously separated from cluster 7 by adding ICOS to the gating strategy, while ConvexGating suggested a two-step strategy with the monocyte markers CD14 and CD16, and previously unused markers CD10 and CD15 (**Fig. 4 C**). ConvexGating corroborated the previous finding where CD16 expression has been identified as a hallmark for activated T cell populations in both the CD4 and CD8 compartment, which was associated with severe outcomes in COVID-19^18^. Adding more hierarchies to the gating strategy strongly improved precision with almost constant F1-score (**Fig.s 4 B - E** leftmost panels). Overall, ConvexGating provided a simplified gating strategy for disease specific T cells.

**Figure 4:**
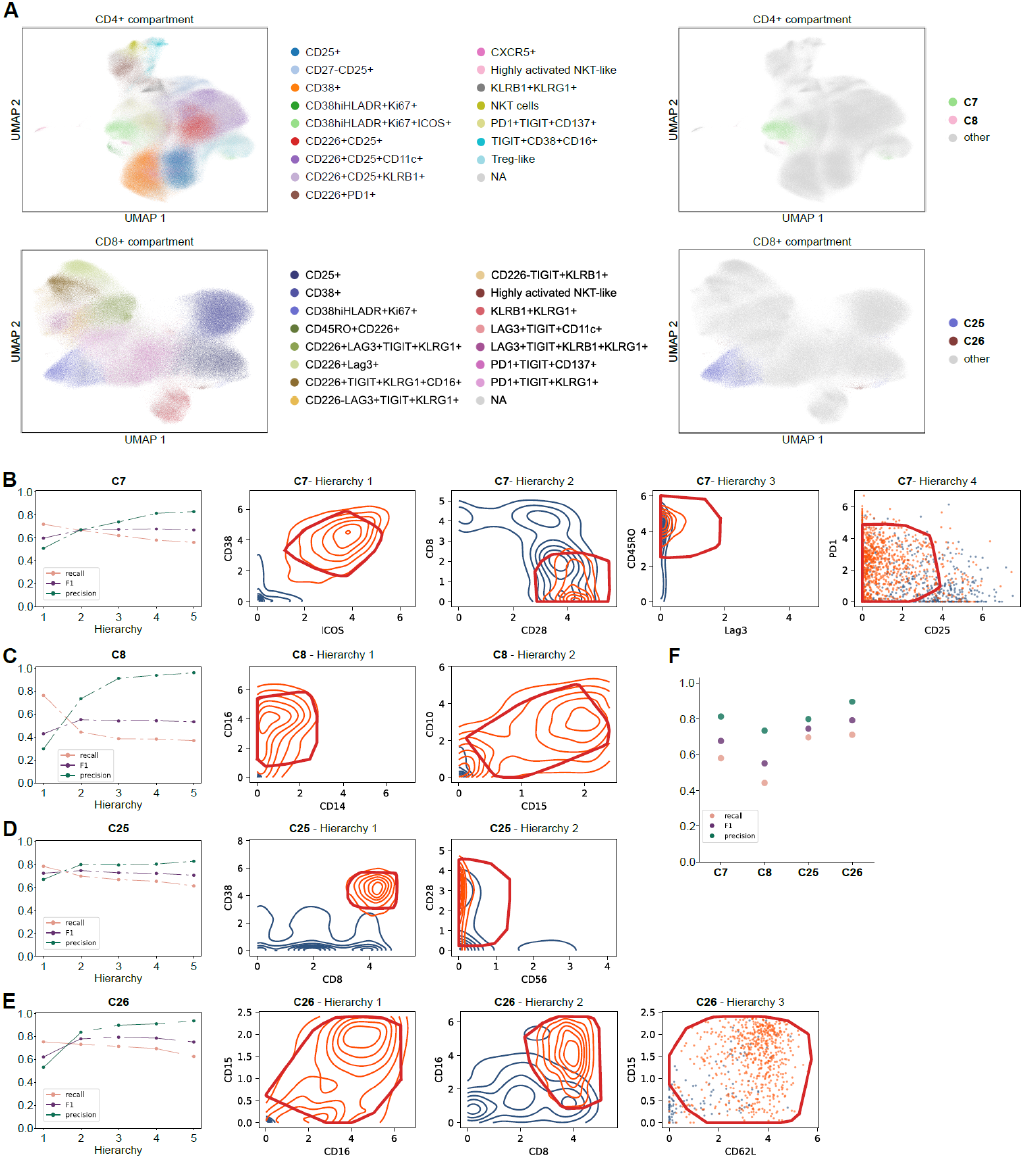
ConvexGating finds gating strategies for unknown populations in cyTOF Covid data^18^. **(A)** UMAP representation of CD4+ T cell compartment (upper panels) and CD8+ T cell compartment (lower panels) with coloring according to subtype annotation (left panels) and clusters (right panels). **(B-E)** Performance scores and gating strategies for cluster 7 - CD4+ T cell compartment **(B)**, cluster 8 - CD4+ T cell compartment **(C)**, cluster C25 - CD8+ T cell compartment **(D)** and cluster C26 - CD8+ T cell compartment **(E)**. Target cells (orange) and non-target cells (blue) are represented via scatter plots or contour plots per target and non-target population. Subsampling was performed to have a ratio of 1:15 between target cells and non-target cells. **(F)** Final performance overview on full cyTOF panel.

### ConvexGating bridges the gap between data modalities

To bridge the gap of sequencing-based cytometry to flow and mass cytometry, we examined ways to leverage large scale panels as created with CITEseq and demonstrate that the inferred marker choice can also be detected in flow cytometry or cyTOF data with a much smaller panel size (**Fig. 5 A**). Here, we show the use of ConvexGating on three independent COVID-19 cohorts measured with CITEseq^34^, cyTOF and FACS^18^ to identify disease specific CD16+ CD4+ and CD8+ T cells, respectively. For the CD16+ T cells in both the CD4 and CD8 compartments measured with cyTOF, we observed a fairly stable performance using overlapping markers of cyTOF and CITEseq (**Fig. 5 B-D**) and cyTOF and FACS, respectively, with the exception of clusters 7 and 25, where the overlapping markers of the cyTOF and FACS panel still allow to identify the population of interest with high recall, but high contamination, too (**Fig. 5 B**). To identify CD16+ T cells in the COVID-19 CITEseq panel, we first denoised the data using totalVI^35^, which integrates both single-cell RNA-seq and protein abundance data, and removes the background noise of the latter (**Supplementary Fig. 11 A-E**). We subsequently identified a fraction of 0.07% CD16+ CD4+ T cells in the CD4 compartment and of 0.11% CD16+ CD8+ T cells in the CD8 compartment, respectively, which are predominantly observed in COVID-19 cases (**Supplementary Fig. 11 E**). Using the full marker panel for each study individually, ConvexGating identified these cell populations with high precision (**Fig. 5 B**). Using only overlapping markers of cyTOF and CITEseq, and flow cytometry and CITEseq, respectively, the performance to identify CD16+ T cells in both compartments remains stable, underscoring ConvexGating’s flexibility to identify target populations from marker subsets (**Fig.s 5 B, E-F)**. For flow cytometry data, we defined a minimal marker panel (CD3, CD4, CD8, CD16 and HLA-DR) to identify our populations of interest. Surprisingly, ConvexGating’s performance even slightly increased on the minimal panel compared to the full marker panel (**Fig.s 5 B, G-H**). In summary, ConvexGating provides interpretable and simple gating strategies for disease specific T cells across single-cell analysis platforms. Moreover, the performance metrics directly indicate what kind of purity level to expect, if those gating strategies are subsequently used for cell sorting and functional experiments on the full panel or on subsets of markers. ConvexGating handles different subsets of marker panels with stable performance. While we provide an export function for gating strategies, transferring them across modalities (FACS, cyTOF and CITEseq) would most likely require manual adjustment. In addition, ConvexGating shows how well populations defined in complex panels can be gated in panels with fewer markers.

**Figure 5:**
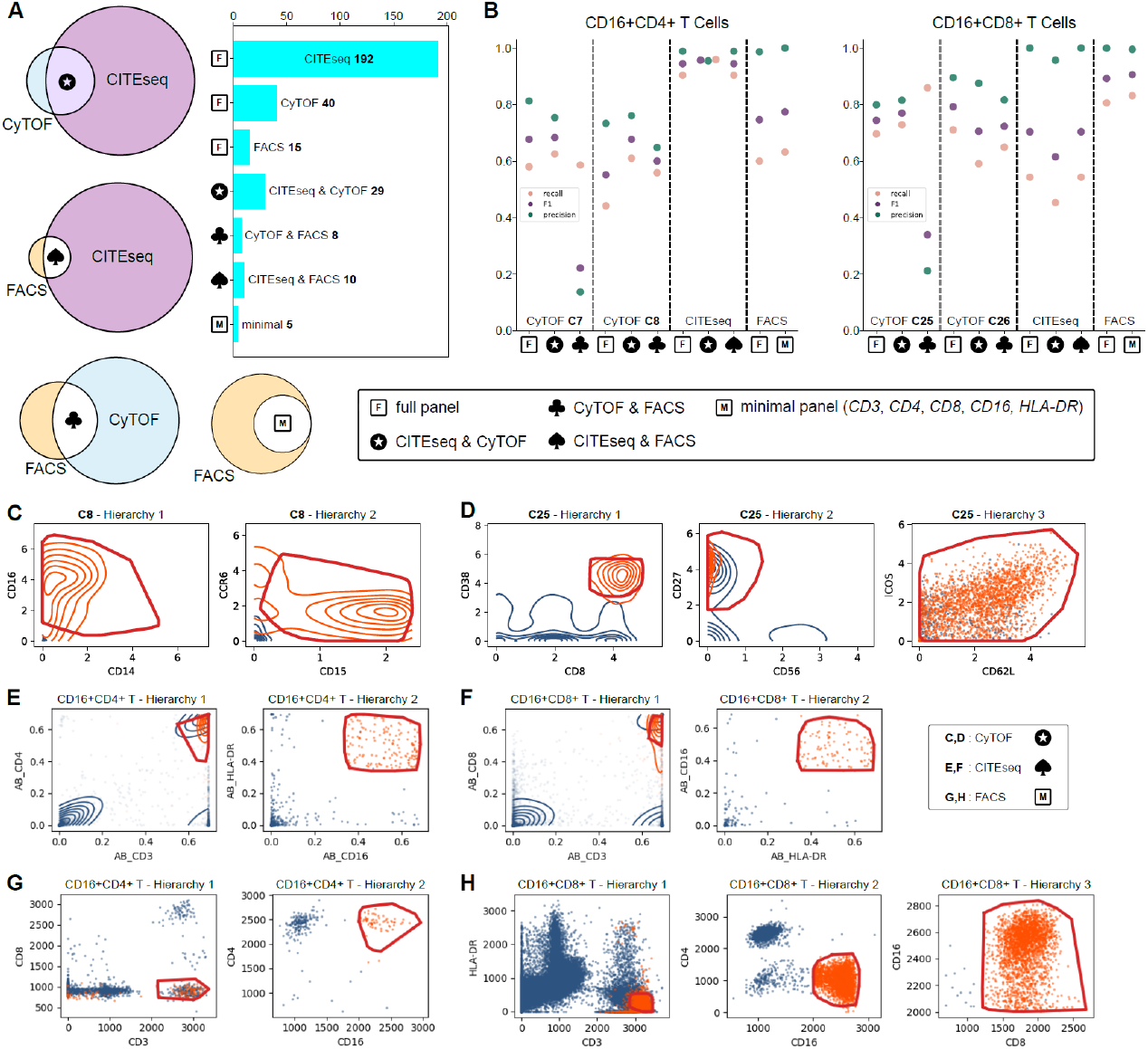
ConvexGating bridges the gap between cyTOF, CITEseq and FACS datasets. **A)** Overlap of cyTOF, CITEseq and FACS antibody panels for independent cohort studies of COVID-19 using peripheral blood. (**B)** Performance overview for identification of CD16+ CD4 T cells (left panel) and CD16+ CD8 T cells (right panel) on full and restricted antibody panels per modality via ConvexGating. **(C-D)** cyTOF: gating strategies for cluster 8 in CD4+ T Cell compartment (**Fig. 4 A**) **(C)** and for cluster 25 in CD8+ T cell compartment (**Fig. 4 A**) **(D)** on joint CITEseq & cyTOF panel. **(E-F)** CITEseq: gating strategy for CD16+CD4+ T cells **(E)** and CD16+CD8+ T cells **(F)** on joint CITEseq & FACS panel. **(G-H)** FACS: gating strategy for CD16+CD4+ T cells **(G)** and CD16+CD8+ T cells **(H)** on minimal panel (CD3, CD4, CD8, CD16). Target cells (orange) and non-target cells (blue) are represented via scatter plots and/or contour plots per target and non-target population. Subsampling was performed to have a ratio of around 1:15 between target cells and non-target cells **(C-H)**.

## Discussion

Machine learning applications have become increasingly popular for flow and mass cytometry data. This development is accelerated by recent technological advances to increase the size of flow cytometry marker panels and the development of joint profiling of transcriptome and epitopes via e.g. CITE-seq. Yet, in flow cytometry, manual gating is still the current Gold standard for both data analysis and sorting of cells of interest.

While an increased flow cytometry panel offers an enhanced characterization of single cells, it also makes the drawbacks of manual gating more apparent. In our study, we observed large inter-rater variation across operators, bias in cell type proportions due to the hierarchical nature of manual gating and inconsistent cell type label assignment especially in the DC compartment (**Fig. 3**). In contrast, clustering leverages the high-dimensional feature space to separate cell types and leads to very similar results as manual gating by an experienced operator (**Fig.s 2, 3**).

As such, data analysis via supervised, unbiased clustering^13^ or advanced ML techniques like automated cell type annotation using normalizing flows^36^ pose an alternative to manual gating. Cell sorting via FACS relies on hierarchical gating instead of clustering methods, such that tools like our proposed ConvexGating provide gating strategies in a data-driven, unbiased manner. Marker choice in ConvexGating depends on separability, such that markers have equal chances of being selected in the gating strategy, regardless of default usage in manual gating or frequent appearance in the literature. ConvexGating offers gating strategies optimized for cell sorting where minimized contamination is a prerequisite. This requires a high degree of precision in the extracted population while still capturing most of the target cells to obtain a comprehensive picture of the population under investigation. ConvexGating puts this into practice by aiming to maximize F1 score with precision implicitly given priority over recall. In terms of F1 score, ConvexGating outperforms the recently released Hypergate gating tool^26^ for fine-grained cell types in different scenarios and for different data modalities (**Supplementary Fig. 9**). Cell type abundance also hampers performance. While Hypergate enriches the target population to a one-to-one ratio, ConvexGating addresses class imbalance with an importance weighting in the loss function (see **methods**). This adaptation accounts largely for the observed class imbalance of target-to-non-target populations, and only for very rare cell types we enriched the target population to a 1:15 ratio to improve performance. ConvexGating provides short, high-precision gating strategies while Hypergate derives high-recall gating strategies with an increased risk for contamination. Consequently, we consider ConvexGating better suited for tasks where a high purity in the extracted population is desired.

It must be noted that repeated mapping of high-dimensional marker expression data into a 2D marker space while striving for short, concise gating strategies is not optimal with respect to classification performance. More flexible ML models, such as non-linear SVM classifiers, separate target populations with higher precision and recall (**Supplementary Fig. 10**). The higher performance comes at the cost of difficult-to-interpret decision boundaries, which lack direct translatability into a gating strategy. Moreover, many of these models likely construct a fragmented classification area in the marker space, which ultimately lacks a concise characterization of the cells under investigation, fails to generalize and limits comparability between samples. Therefore, ConvexGating offers a trade-off between performance and interpretability. Convex gate shapes induce contiguous, interpretable gated areas in 2D marker space while at the same time enabling more target population-adaptive and flexible gates compared to rectangular shapes or marker thresholding^20,21,26,27^.

Due to its data-driven nature, ConvexGating supports the exploration and characterization of newly discovered or rare cell populations. The used markers indicate how the cell population can be efficiently extracted, regardless of whether the cell populations were originally identified via clustering or manual gating. The derived gating strategies then in turn allow drawing conclusions about the underlying biology of the cell populations under investigation (**Fig.s 4, 5**) and allow for optimal panel design. For high-dimensional human bone marrow data measured with cyTOF, ConvexGating provides high-precision strategies for both broad and fine-grained cell type definitions (**Supplementary Fig. 8**). Notably, rare T cell subsets require more markers for precise gating. At the same time, ConvexGating uses markers that provide the best separability for a population of interest. In the disease context, ConvexGating is capable of deriving gating strategies for unknown cell subpopulations, e.g. the COVID-specific T cell subpopulations in the CD4+ and CD8+ T cell compartments (**Fig. 4**). Here, ConvexGating helps with marker prioritization as a number of markers were found to describe the unknown population, and we showed that a minimal panel of four or five markers, respectively, is already sufficient to describe these cells. Large panels, as in cyTOF, spectral flow or CITEseq experiments, provide a comprehensive picture of the cells under study, but translation into smaller panels while preserving performance might not be straightforward. In this scenario, ConvexGating can be used for optimal antibody choice and panel design as we can determine the performance of a smaller panel compared to larger panels computationally (**Fig. 5**). Such an approach fosters the standardization of marker choice from extensive cellular signatures to effective sorting strategies. We demonstrated how ConvexGating can be used to create gating strategies for perturbed populations across multiple studies and modalities. However, a direct translation of a gating strategy would still require manual adjustment. We consider the translation of the inferred gating strategies from CITEseq or cyTOF data to a flow cytometry or spectral flow experiments as a separate task since the data distribution in each modality is quite different, such that a reliable transfer of a gating strategy requires an invertible mapping into the same feature space. Finding such potentially nonlinear mappings is challenging because a reliable mapping of the same cells must be established as well as a meaningful mapping of the convex gates from the integrated data space back into the original data space that is used for cell sorting.

In conclusion, ConvexGating links machine-learning based data analysis with cell identification by deriving gating strategies mimicking expert manual gating in an automated fashion. While some markers can be used interchangeably, our approach contributes to the efforts in standardization of cell type labeling and subsequent sorting. Our developed Python package integrates seamlessly with other popular single-cell analysis frameworks in Python to facilitate ML for flow and mass cytometry data. We envision that the explainability of the ConvexGating model and its output fosters an enhanced acceptance of our approach in biological and medical research as well as drug discovery, leading to more efficient analysis pipelines that can adapt to expanding parameter spaces. In the future, we believe that machine-learning tools like ConvexGating could lay the foundation for potential sample-specific automated live adjustments on cell sorters.

## Methods

### Staining of whole blood of healthy donors (DC panel)

The antibody panel was designed to analyze monocytes and cDC2 subsets^31^. 100 µl of heparinized venous blood from three apparently healthy donors was used and subjected to erythrocyte lysis using Q-Prep (Coulter). After centrifugation, the cell pellet was resuspended in 100 µl staining-buffer (PBS + 2% FCS + 2 mM EDTA) and cells were incubated with CD1c-PE, CD5-BV711, CD14-FITC, CD16-APC, HLA-DR-PE-Cy7, CD19-AF700, and CD20-AF700 for 20 min in the dark on ice. After washing, the cells were resuspended in a staining-buffer and analyzed using a Coulter cyTOFLEX LX flow cytometer. A compensation matrix was generated using single stained Versa Comp compensation beads (Coulter) and manually adjusted. Fcs-files were analyzed using FlowJo v10.8. After gating on singlets plus HLA-DR+ non-B cells, monocyte subsets were defined based on expression of CD14 and CD16, and cDC2 subpopulations were identified using CD1c, CD14, and CD5^31^ (**Supplementary Fig. 1**).

## Data analysis

In the following, we describe how we carry out the preprocessing steps with pytometry^14^, and scanpy^15^. Let *X*^*orig*^*ϵ R*^*NxM*^ be our data matrix that contains original measurement values stored as one FCS file per sample. *N* denotes the number of cells and *M* denotes the number of markers. The data processing steps vary in every dataset and are described in the following sections. After our preprocessing steps, we refer to our data matrix as *X*^*data*^*ϵ R*^*NxM*^. Unless stated otherwise, all analyses were carried out in Python v. 3.8 with scanpy v. 1.8.1, anndata v. 0.8.0 and pytometry v. 0.1.2.

### Flow cytometry data of whole blood of healthy donors (DC panel)

The dataset of whole blood stains encompasses three healthy donors stained with six markers (see **DC panel** above). We load all data in Python (v. 3.8) using the pytometry package^14^ (v. 0.1.3). We convert the FCS files into the anndata^16^ format (*pytometry*.*io*.*read_fcs*). Next, we perform compensation to correct for potential fluorescent spillover across channels, i.e. by multiplying the inverse of the spillover matrix to the data matrix. Then, we employ the bi-exponential transformation for normalization of flow data. We then subset the data using the manual singlet gate and perform Leiden clustering^12^ of the k-nearest neighbor graph directly on the data (*scanpy*.*pp*.*neighbors*)^15^ to group cells and annotate the clusters based on characteristic marker intensities (scanpy v. 1.8.2). In particular, we employ a split-and-merge strategy to annotate the cells at different levels of resolution. In the first round, we separate B cells, monocytes and DCs from all other cells, which we term “not annotated” (**Supplementary Fig. 2**). In the next step, we subset to monocytes and DCs, and re-computed the k-nearest neighbor graph and Leiden clustering to identify intermediate and non-classical monocytes (**Supplementary Fig. 3**). We reiterate this procedure for the DCs to identify DC2 subsets and to separate the remaining classical monocytes (**Supplementary Fig. 4**). We use UMAP to represent the data with low dimensional visualizations (*scanpy*.*tl*.*umap*). In the final step, we merge the resulting cell type annotation into broad categories (cell type level 1) and refined categories (cell type level 2). To check the coherence of manual gating (**Supplementary Fig. 1**) and clustering, we visualize the final annotation as 2D scatter plots (**Supplementary Fig. 5**).

We performed ConvexGating with standard parameters per donor and per cell type on a subset of 50,000 cells (*scanpy*.*pp*.*subsample*) for both cell type level 1 and cell type level 2 (**Supplementary Fig. 6**).

### Flow cytometry data of frozen PBMCs of healthy donors (Knoll et al.)

Flow cytometry data were originally collected by Knoll et al.^32^ using a 27 marker panel (see Supplementary Table S1 in Knoll et al.) and data were shared with us by the authors. In this study, we use five samples collected from healthy donors. We load all data in Python (v. 3.8) using the pytometry package^14^ (v. 0.1.2). We convert the FCS files into the anndata^16^ format (*pytometry*.*io*.*read_fcs*). Next, we perform compensation to correct for potential fluorescent spillover across channels, i.e. by multiplying the inverse of the spillover matrix to the data matrix. We then remove doublets from the data via FSC-A/SSC-A and FSC-A/FSC-H channels. We then normalize with the bi-exponential transformation similarly to the original analysis with FlowJo (BD, v. 10.7.1).

We then perform Leiden clustering^12^ of the k-nearest neighbor graph directly on the data (*scanpy*.*pp*.*neighbors*)^15^ to group cells and annotate the clusters based on characteristic marker intensities with a similar split-and-merge strategy as described above (**Supplementary Fig. 7**) using scanpy v. 1.9.1.

### cyTOF data of human bone marrow (Oetjen et al.)

Leveraging previous work on the pytometry package^14^, we downloaded the publicly available flow and mass cytometry data of healthy human bone marrow donors^33^ from FlowRepository.org (accession codes: FR-FCM-ZYQ9, FR-FCM-ZYQB). We normalized the data with arcsinh-transformation and cofactor 5. We then use previously created cell type annotation based on Leiden clustering and marker intensities. The detailed analysis can be found at https://pytometry.readthedocs.io/en/latest/examples/.

We applied ConvexGating to retrieve gating strategies for cell populations in cyTOF data of human bone marrow. Gating strategies were separately learned on eight samples and four annotation levels (level 2 - level 5) (**Supplementary Fig. 8**). In level 2, NK cells and T cells were annotated. With increasing annotation level, T cells were further subtyped (**Supplementary Fig. 8 A**). We subsampled to 50,000 cells (*scanpy*.*pp*.*subsample* in scanpy v. 1.8.1) before applying ConvexGating with standard parameters. For *CD8+ TRM T cell* population in *sample B*, we ensured a minimum number of 100 target cells after subsampling. We evaluated the performance of the gating strategies as described above. We further compared the performance of the full gating strategy with the performance of a reduced gating strategy that only relies on the two initial hierarchies. For all inferred gating strategies, we determined the optimal number of hierarchies and visualized the distribution per cell type annotation level (**Supplementary Fig. 8 B**).

### cyTOF data of T cells of COVID-19 patients

We downloaded the normalized and annotated cyTOF data set of T cells of COVID-19 patients^18^ from https://zenodo.org/record/5771937/files/data_Tcells_annotated.csv.gz. We then converted the data frame into an anndata object, separating marker intensities and metadata. Next, we selected the previously identified CD16+ T cells populations in the CD4 and CD8 compartment, which were annotated as clusters 7 and 8, and clusters 25 and 26, respectively. For each target cluster, we subsampled the anndata object to have an equal proportion of 6.25% target cells and 93.75% non-target cells (proportion 1:15). Furthermore, for cluster 7 and cluster 25 we subsampled the corresponding anndata object to 50,000 cells (*scanpy*.*pp*.*subsample*). We then performed ConvexGating with standard parameters.

### Flow cytometry data of T cells of COVID-19 patients

We obtained the compensated flow cytometry data set of T cells of COVID-19 patients^18^ as an inhouse data set. We convert the FCS files into the anndata^16^ format (*pytometry*.*io*.*read_fcs*). We then normalize the data using the bi-exponential transformation as previously used for analysis with FlowJo (BD, v. 10.7.1). Within the CD4 and CD8 T cell gates, to identify CD16+ T cells, we used a normalized CD16 marker intensity above 2,000 similar to the CD16 threshold for neutrophils. The fraction of CD16+ CD4+ T cells is 0.048% of total CD4+ T cells and 2.54% of total CD8+ T cells.

For each target population, we subsampled the anndata object to have an equal proportion of 6.25% target cells and 93.75% non-target cells (proportion 1:15). Furthermore, for CD16+ CD8+ T cells, we subsampled the corresponding anndata object to 50,000 cells (*scanpy*.*pp*.*subsample*). We then performed ConvexGating with standard parameters.

### CITEseq data of the immune cell compartment in COVID-19 patients

We downloaded the annotated CITEseq data set of the immune cell compartment of COVID-19 patients and healthy donors^34^ from https://www.covid19cellatlas.org/. The CITEseq dataset consists of 624,325 cells with 24,737 expressed genes and 192 markers, respectively. In order to define CD16+ T cells, we selected the “raw” count data from the data set. We denoised the CITEseq data using totalVI^35^ to account for the background signal of CITEseq data. For cell type definition, we rely on the previously published cell type definition for CD4+ and CD8+ T cells (**Supplementary Fig. 11 A**). We denote HLA-DR+ CD16+ T cells as T cells with a foreground probability of both HLA-DR and CD16 higher than 0.4 and being clustered with the CD4 and CD8 T cell compartment, respectively (**Supplementary Fig. 11 B-E**). As part of the preprocessing pipeline, we utilize log+1 scaling for normalization of the protein expression data. Furthermore, we subsample the data for each cell type of interest, namely CD4 and CD8 T cells, to ensure a proportion of 6.25% target cells and 93.75% non-target cells before applying ConvexGating with standard parameters.

### Manual Gating comparison

To assess the inter-rater variability of manual gating across operators, we devised the following manual gating task to be carried out by members of the lab. The test dataset was originally collected by Knoll et al.^32^ and consisted of PBMCs samples from five healthy human donors measured on a BD Symphony A5 instrument, encompassing 27 markers (**Supplementary Fig. 7 A**), which covers the markers for all major immune cell populations. The dataset was compensated, but not filtered for debris. Every participant is trained in the use of a flow cytometer and in the analysis of immune cell populations using FlowJo (BD, v. 10.7.1). We obtained 11 submissions from 7 participants, where one participant submitted 3 different gating strategies. As “Gold standard”, we defined the cell type annotation of the original study. All participants were asked to gate the following populations:

B cells, T cells (with the subtypes CD4+, CD8+ and NKT cells), NK cells, monocytes (with subtypes classical, non-classical, intermediate), and DCs (with the subtypes pre-DC, pDC and cDCs). No further advice as to which gating strategy to use was given. Notably, the task was designed to cover abundant populations as well as more rare and difficult to gate ones.

All participants submitted their manual gating results in FlowJo as FlowJo workspace (wsp) file. For analysis, we converted all gating results into a one-hot encoded data frame using CytoML v. 3.12 and flowWorkspace v. 3.13 in R v. 4.0.3 and exported the cell annotation data frames then as csv files for further analysis with pandas v. 1.3.5 and scanpy v. 1.9.1 in Python v. 3.8. We harmonized cell type labels into three levels: The “valid cell” level marks the first pass QC, where debris is removed from further analysis. Next, we defined the first level of cell type annotation as the cell types B cells, T cells, NK cells, monocytes, and DCs. The second level of cell type annotation encompassed in addition the subtypes of T cells, monocytes, and DCs, respectively. Cells that were not classified or failed QC were labeled as “not annotated”. In addition to manual gating, we analyzed the data using Leiden clustering in scanpy and added this annotation as “clustering” to the comparison (see **data analysis**).

We examined the concordance of cell type labeling using manual gating and clustering annotation with the Jaccard index, normalized mutual information, and inverse Simpson index (ISI), and determined the cell type compositions for all submissions. In addition to the Gold standard, we computed the most often assigned label as “consensus label” as the most likely label of a cell to account for potential bias in the Gold standard. Unless stated otherwise, we compared cell type annotations to the Gold standard.

### Jaccard index

The Jaccard index^37^ quantifies the similarity of two set *S*_1_ and *S*_2_ by considering its intersection over union.

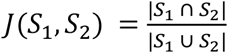

It follows that 0 ≤ *J*(*S*_1_, *S*_2_) ≤ 1. High values for *J* indicate high similarity of *S*_1_ and *S*_2_.

### Normalized Mutual Information (NMI)

Mutual information (MI) between two label assignments of the same data quantifies the degree of similarity between the labels^38^. The NMI further maps the MI to the range [*0,1*] where higher NMI values indicate higher similarity between two sets of labels.

### Inverse Simpson Index (ISI)

Inverse Simpson Index (ISI) is a measure for the effective number of categories present in a population. It is based on the Simpson Index (SI)^39^, which measures the diversity within a population. For a population *P* with *N* total elements, *C* different categories and *n*_*c*_ elements per category, SI reads as

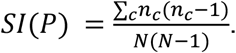

We have that 0 < *SI*(*P*) ≤ 1. If *SI*(*P*) = 1, all elements stem from one category while for *SI*(*P*) = 0 each element is assigned to its own category. The higher SI, the lower the diversity within population *P*. The ISI score is then obtained by taking the inverse of SI

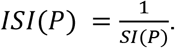

It follows that 1 < *ISI*(*P*) ≤ *C*. ISI reflects the effective number of categories in *P*. To determine the most likely label of a cell, we used the most often assigned label as “consensus label” to account for potential bias in the Gold standard. The lower the ISI for a cell type reflects a more consistent labeling across operators.

### Benchmark Gating Strategies

We conducted a comprehensive benchmark analysis of ConvexGating against the state-of-the-art automated gating tool Hypergate^26^. Hypergate fits a high-dimensional rectangle to separate target cells from non-target cells. Every projection of the high-dimensional rectangle into 2D space is again a rectangle, which is then used to derive a marker set as a gating strategy. Furthermore, we benchmarked ConvexGating against two supervised binary classification algorithms, namely a linear support vector machine (SVM) classifier and a radial basis function (rbf) kernel SVM^38^. The linear SVM directly searches for one separating hyperplane in M dimensional marker space that optimally distinguishes target cells from non-target cells. The rbf SVM is based on a non-linear rbf kernel transformation that maps the input data into a vector space with enhanced separability of targets and non-targets. However, the increased separability induced by a non-linear mapping comes at the cost of interpretability. In contrast to Hypergate and ConvexGating, the output of the SVM classifiers does not directly translate into a gating strategy. Both SVM classifiers were implemented with the aid of scikit-learn python package v. 1.1.2^38^.

For executing hypergate, we made use of the most recent hypergate R package version 0.8.3 (https://github.com/ebecht/hypergate) in R v. 4.0.3. Following the recommendation of the package we sampled target and non-target population to 1:1 ratio with 2000 cells each. The resulting gates were then applied to all cells for performance evaluation. Since ConvexGating and both SVM classifiers do not require a 1:1 ratio of target and non-target cells, we sampled 50,000 cells.

The benchmark analysis was conducted for two different flow cytometry data sets (DC panel, large PBMC panel) with two annotation levels each (level 1, level 2). Furthermore, the human bone marrow cyTOF dataset (Oetjen et al.^33^ see **data analysis**), which consists of four annotation levels (level 2, level 3, level 4, level 5), was included in the benchmark analysis. As performance measure for this benchmark analysis we referred to precision, recall and *F*_1_ score (see **performance evaluation**).

### Performance evaluation

The goodness of a gating strategy is evaluated based on recall, precision and *F*_1_ score. Recall quantifies the portion of target cells (positives) identified by the gating strategy. Precision refers to the fraction of target cells inside the gated population. A high recall value indicates that most target cells are captured by the gating strategy while a high precision value indicates purity in the gated population with target cells predominating over non-target cells (negatives).

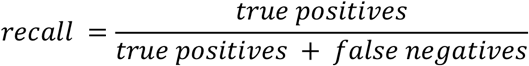

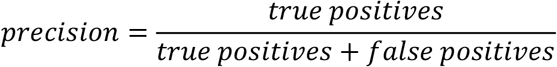

*F*_1_ is the harmonic mean of precision and recall. For a gating strategy, a high *F*_1_ indicates both a high identification rate of target cells and purity in the gated population.

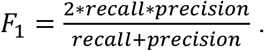

### ConvexGating Algorithm

Let *X*^*data*^*ϵ R*^*NxM*^ be our data matrix where *N* denotes the number of cells and *M* denotes the number of markers. Optionally, we add a small amount of random noise from a uniform distribution over [-5*e*^−5^, 5*e*^−5^) per entry of *X*^*data*^ to ensure robust internal processes within our algorithm, particularly in cases where many cells exhibit zero expression for a specific feature. Cells are assumed to be labeled, *i. e*. assigned to one of *C* cell populations (clusters). Our model is capable of learning a gating strategy for extracting specific cell populations from the remaining cells in an interpretable way, mimicking the behavior of human experts performing manual gating. Cells of the desired population are referred to as targets. All other cells are referred to as non-targets.

### Deriving Gating Strategies

We define *T* as the maximum number of hierarchies in the gating strategy while *F* denotes the set of available markers. Per hierarchy we have a target population *P*^*t*^ and a non-target population *P*^*nt*^. Finding an appropriate gate per hierarchy is a three-step procedure, which we describe in the following.

#### Step 1. Selecting Marker Combination for 2D Marker Space

For *P*^*t*^ and *P*^*nt*^, we calculate the 1*st*, the 50*th* and the 99*th* percentile along each marker *f ϵ F*. These statistical quantities are used as a heuristic for the distribution of *P*^*t*^ and *P*^*nt*^ along marker *f*. We select those two markers that promise the largest difference in distribution for *P*^*t*^.

For each marker*f ϵF*, we define

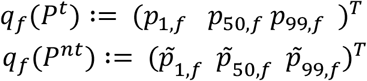

where *p*_*u,f*_ denotes the *uth* percentile of *P*^*t*^ along marker *f* and *p* _*u,f*_ the *uth* percentile of *P*^*nt*^ along marker *f*. We then calculate for all *f ϵ F* the heuristic

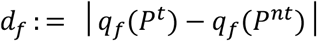

and select those two markers with the highest heuristic value for the corresponding hierarchy.

#### Step 2. Learning Gate Location

After selection of markers *m*_1_and *m*_2_, we learn the optimal location of gate *G* in 2*D* marker space by minimizing a problem-specific *loss* via *stochastic gradient descent*.

### Gate specification

We define a gate in 2*D* marker space as the intersection of *K* halfspaces *H*^+^ in 2*D*. Ahalfspace *H*^+^ is determined by a 2*D* hyperplane *C* with normal vector *w* ∈ *R*^2^ and bias *b* ∈ *R* which reads as

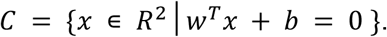

The corresponding halfspace *H*^+^ takes the form

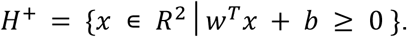

Our convex gate *G* is then specified as

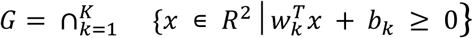

where *w*_*k*_ ∈ *R*^2^ denotes the *normal vector* and *b*_*k*_ ∈ *R* the *bias* of the *kth* hyperplane.

#### Loss function

Our loss *L* quantifies how well gate *G* captures target cells and separates them from non-target cells. We apply a weighted version of *binary cross-entropy loss*. For marker expression input *x* ∈ *R*^2^ with label *y* ∈ {0,1} and weight parameter *α*, our loss *L*_*α*_ computes as follows. For all hyperplanes *k* = 1, …, *K* we calculate

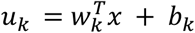

which puts the location of input *x* into relation with the *kth* hyperplane. In case *x ϵ G*, we have for all *k* = 1, …, *K* that *u*_*k*_ ≥ 0. Next, we calculate the predicted label

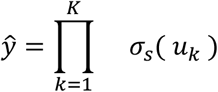

where *σ*_*s*_ denotes the parameterized sigmoid which is defined as

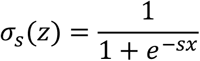

for an input *z ϵ R* and parameter *s ϵ R* (default: *s*=40). For sufficiently large *s ϵ R*, the parameterized sigmoid *σ*_*s*_ approximates the indicator function. Finally, our loss *L*_*α*_ reads as

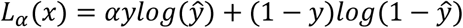

where *α* acts as a weight parameter adjusting the influence of target cells to the loss. Let *n*_*target*_ > 0 denote the number of target cells and *n* _*non*−*target*_ the number of non-target cells.

We set

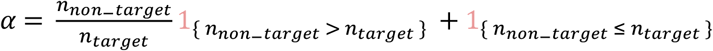

where 1_{}_ denotes the indicator function.

### Regularized loss

We add two problem-specific regularization terms *P*_1_ and *P*_2_ to the loss *L*_*α*_ that encourage tight gates around the target population. The regularized loss is referred to as 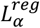 (see **Supplementary Note 3** for details):

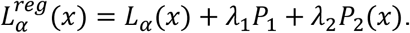

The weight parameters *λ*_1_ and *λ*_2_ control how much each regularization term affects the loss.

### Loss minimization

We obtain the optimal gate parameters *w*_1_, …, *w*_*K*_ and *b*_1_, …, *b*_*K*_ via stochastic gradient descent. We start by a random parameter initialization. Then, we repeatedly take a subset *S* ⊆ *P*^*t*^ ∪ *P*^*nt*^, compute 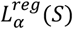 and backpropagate the loss to the gate parameters. We then update for *k* = 1, …, *K* the normal vectors and biases based on the derivatives

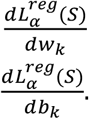

To control for the randomness induced by random parameter initialization, we fix four (default) normal vectors based on the principal components of the target population *P*^*t*^, thus stabilizing training.

### Adaptive grid search

Finding an optimal gate *G* requires a careful selection of the weight parameters *λ*_1_ and *λ*_2_ that control the influence of the penalty terms *P*_1_ and *P*_2_ in the regularized loss 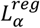 (see **Supplementary Note 3**). We perform an adaptive grid search to find those weight parameters *λ*_1_ and *λ*_2_ that lead to the best gate *G* with respect to separation of target cells and non-target cells (performance measure: *F*_1_ (default) or recall). To reduce the computational complexity, we assign equal weight to both penalty terms *P*_1_ and *P*_2_, thus setting *λ*_1_ = *λ*_2_.

To determine *λ*_1_, *λ*_2_, we define an initial range (default: [0, 2, 4, 8, 10]) of possible weight parameters that is iteratively updated based on the two best-performing weight parameters *h*_1_ and *h*_2_ in the current range of values. Let *grid*_*divisor*_ > 0 and define

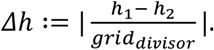

The updated search range then starts either at *min*{*h*_1_, *h*_2_} − *Δh* or at *min*{*h*_1_, *h*_2_} + *Δh* and extends until *max*(*h*_1_, *h*_2_) + *Δh* with uniform steps of size *Δh* in-between. The number of search range updates can be chosen manually (default: 2). Eventually, the gate leading to best performance during the adaptive grid search is chosen as gate *G*.

#### Step 3. Gate optimization via convex hull

Let 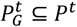 denote the subset of the target population that contains all target cells inside gate *G* after step 2. As final gate *G*^*final*^ we take the convex hull of 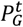, that is

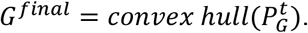

*G*^*final*^ encompasses the same target cells as *G*, but is only as large as necessary while still being convex. Non-targets contained in *G* drop out if they lie outside the convex hull of 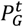.

## Evaluation

The goodness of a gating strategy is evaluated based on recall, precision and *F*_1_ (see section Performance evaluation). Our model is encouraged to optimize *F*_1_ with an emphasis on precision (default) as we aim to construct gates with high cell type purity, such that our gating strategies can be applied for cell sorting and subsequent functional testing of the cells.

## Code and data availability

ConvexGating is implemented in Python, available on PyPI and on GitHub https://github.com/buettnerlab/convexgating. Documentation and tutorials can be found on https://convexgating.readthedocs.io/en/latest/index.html. All scripts and notebooks used to create the figures are available on https://github.com/buettnerlab/reproducibility_convex_gating.

## Author contributions

M.B. conceived the study with the support from J.L.S. and F.J.T.

T.P.H. and E.N. performed experiments and analyzed data.

V.D.F., L.B., M.B., M.D.B., K.M., A.F., C.L.H., C.C., M.H.-S. and D.H. performed data analysis.

V.D.F., L.B. and M.B. wrote the code.

M.B., L.B., M.D.B., M.S., J.L.S. and F.J.T. supervised the work.

V.D.F., L.B., M.D.B. and M.B. wrote the manuscript with the help of all co-authors. All authors read and corrected the final manuscript.

Acknowledgments

V.D.F. was funded by the Federal Ministry of Education and Research of Germany (BMBF; 01IS18026B) and by Sächsische Staatsministerium für Wissenschaft, Kultur und Tourismus in the programme Center of Excellence for AI-research, „Center for Scalable Data Analytics and Artificial Intelligence Dresden/Leipzig”, project identification number: ScaDS.AI.

M.D.B. is supported by the excellence cluster ImmunoSensation2 (EXC 2151, #390873048); the DFG via IRTG2168 (#272482170), SFB1454 (#432325352); the EU-funded project NEUROCOV receiving funding from the RIA HORIZON Research and Innovation under GA No. 101057775; the Else-Kröner-Fresenius Foundation (2018_A158).

## Declaration of generative AI and AI-assisted technologies in the writing process

During the preparation of this work the authors used ChatGPT/OpenAI for assistance in improving the readability and quality of the English language in this manuscript. After using this tool/service, the authors reviewed and edited the content as needed and take full responsibility for the content of the publication.

