## Supplementary Information for "ConvexGating infers gating strategies from clusters in single cell cytometry data"

### Supplementary Note 1

**ConvexGating finds gating strategies for low and high resolution cell types in healthy human bone marrow mass cytometry data**

To assess the applicability of ConvexGating to high-dimensional mass cytometry data for optimal panel design, we applied ConvexGating to extract cell populations in eight healthy human bone marrow samples measured by mass cytometry[33] with 34 markers, which focused on T cell subtyping. We used four different annotation levels as described previously[14] (**Supplementary Fig. 8 A**). In the full strategy, we allowed a maximum number of five gating hierarchies. We compared the performance of the full strategy (**Supplementary Fig. 8 B** left panel) to the performance of a reduced strategy (**Supplementary Fig. 8 B** right panel) that only relied on the initial two gating hierarchies of the full strategy. We observed a decrease in F1 score, recall and precision with increasing cell type annotation level. This aligns with the circumstance that certain T cell subpopulations such as *Double negative T cells*, *Double positive T cells* or *CD8+ CM T cells* are much more challenging to capture with a gating strategy. Specifically, mean recall (full strategy) dropped from 0.814 in level 2 to 0.548 in level 5, while mean precision (full strategy) gradually decreased with increasing annotation level (0.994 for annotation level 2 vs. 0.833 for annotation level 5). This showcases how ConvexGating aims at avoiding contamination with undesired cell types in the gated population prioritizing precision over recall. Mean F1 score (full strategy) dropped from 0.889 in annotation level 2 to 0.635 in annotation level 5. The performance of the reduced strategy closely resembled the performance of the full strategy for annotation level 2: With increasing annotation level, finer subpopulations were harder to extract when solely relying on four markers in the reduced strategy. This resulted in significantly lower mean precision scores for the reduced strategy in level 3 (0.729), level 4 (0.663) and level 5 (0.551) compared to the full strategy in level 3 (0.904), level 4 (0.894) and level 5 (0.833). On average, the number of gating hierarchies (full strategy) increased with higher cell type annotation level as finer subpopulations required more steps for extraction (**Supplementary Fig. 8 C**). Next, we inspected the number of markers that ConvexGating required for extracting all cell populations across all eight samples per cell type annotation level (**Supplementary Fig. 8 D**). In level 2, ConvexGating used 12 out of 34 possible markers for retrieving T cells and NK cells across eight different samples, where the full and the reduced strategies were based on the identical marker panel. The number of used markers increased with higher cell type annotation level. Gating strategies for cell types in level 3 were based on 25 markers, in level 4 on 27 markers and in level 5 on 31 markers (full strategy). The reduced strategies consumed fewer markers - 17 for level 3, 23 for level 4 and 25 in level 5 - at the cost of lower precision compared to the full strategies. Notably, the number of markers used by ConvexGating is higher than the number of markers used in the manual gating strategy for this dataset. To further examine the marker choice of ConvexGating, we compared how often each marker was used in the full and the reduced strategies for annotation level 5 (**Supplementary Fig. 8 E**). High marker appearance frequency in the reduced strategies indicates a prominent role in primary, broad identification of a subpopulation. Markers that were frequently used in later hierarchies play an important role in the fine-tuning steps. We observed that T cell markers CD4, CD8 and CD197 (CCR7) as well as monocyte marker CD16 appeared most frequently in both full and reduced gating strategies for annotation level 5. Notably, T cell marker CD3 was mostly used in the initial two hierarchies (37 out 45 total appearances) which suggests a key role in the first filtering steps of T cell subpopulations. In contrast, marker CD194 (CCR4) was frequently used in the full strategies (34 times) but barely in the initial two hierarchies (5 times) which suggests an important role in the fine-tuning steps. Therefore, ConvexGating offers the possibility of panel size reduction while maintaining high precision. Markers with very few appearances in the full strategies or the reduced strategies, respectively, could be removed with very little gating performance sacrifice. We showcase the performance overview (**Supplementary Fig. 8 H**) and the complete gating strategies for level 2 T cells from sample A (**Supplementary Fig. 8 F**) and level 5 CD4+ TEMRA cells from sample A (**Supplementary Fig. 8 G**) found by ConvexGating. The optimal gating strategy for level 2 T cells could be found in two hierarchies. Notably, the precision of the gating strategy for T cells increased marginally in the second hierarchy (**Supplementary Fig. 8 H** left panel). Therefore, the T cell gating strategy could be even simplified to a single step of gating CD3 vs CD2. For level 5 CD4+ TEMRA cells, ConvexGating required five gating hierarchies for an optimal strategy, where it starts already with the markers CD4 and CD57, instead of CD3 to label CD4+ T cells (**Supplementary Fig. 8 G**). This demonstrates an important difference of ConvexGating to traditional gating strategies: While manual gating infers a full hierarchy from broadly to finely resolved cell type definitions, ConvexGating directly infers a gating strategy for a target population, regardless of potential parent populations.

#### Supplementary Note 2

**ConvexGating is better suited for high resolution cell types compared to Hypergate**

We sought to compare the performance of ConvexGating against the most recent gating tool Hypergate[26] (**Supplementary Fig. 9**) and the SVM classifier (with linear and RBF kernel, see **methods** and **Supplementary Fig. 10**). We benchmarked all tools on three test scenarios: the DC panel with two cell type annotation levels (**Fig. 2**), the large PBMC panel with two cell type annotation levels (**Fig. 3**) and the T cell cyTOF panel of human bone marrow data with four annotation levels[14,33] (**Supplementary Fig. 8** and **Supplementary Note 1** for details). For all samples and cell type annotation, we evaluated using F1 score, precision and recall (see **metrics**). We observed a slightly higher mean F1 score for Hypergate (0.906) compared to ConvexGating (0.887) for annotation level 1 in the DC panel (**Supplementary Fig. 9 A** upper left panel). The finer cell subpopulations (annotation level 2) were more challenging to capture with a gating strategy. Here, ConvexGating clearly outperformed Hypergate for nearly all cell subpopulations in terms of F1 score (**Supplementary Fig. 9 A** upper right panel). The higher F1 scores were based on the substantially higher precision of the gating strategies derived by ConvexGating (**Supplementary Fig. 9 A** mid panel). The extracted populations were less cross-contaminated with undesired non-target cells when relying on the gating strategies learned by ConvexGating. Gating strategies from Hypergate exhibited higher recall compared to ConvexGating (**Supplementary Fig. 9 A** lower panel), however, the higher recall did not suffice to equalize the substantially lower precision compared to gating strategies learned by ConvexGating. For broad cell type definitions in annotation level 1 of the large PBMC panel, mean F1 score for gating strategies derived by Hypergate (0.914) was above the mean F1 score for gating strategies derived by ConvexGating (0.858) (**Supplementary Fig. 9 B** upper left panel). However, for finer subpopulations in annotation level 2, ConvexGating outperformed Hypergate with regards to mean F1 score (0.786 vs. 0.743) (**Supplementary Fig. 9 B** upper right panel). Again, gating strategies derived by ConvexGating prioritized precision (**Supplementary Fig. 9 B** mid panel) whereas gating strategies derived by Hypergate exhibited higher recall (**Supplementary Fig. 9 B** lower panel). We observed a similar pattern on the cyTOF panel of human bone marrow data (**Supplementary Fig. 9 C**). For annotation level 2 with low-resolution NK Cell and T cell labels, we observed a slightly higher mean F1 score for gating strategies learned by Hypergate (0.922) compared to gating strategies learned by ConvexGating (0.889). The increasing annotation level subtyped T cells leading to smaller cell subpopulations, which were harder to extract by a gating strategy. The reason for this is that the influence of a single cell outside the gate on recall is more fatal in small populations than in large populations. However, gates that are too large lead to severe contamination (low precision) when a high number of non-target cells are present, making gating particularly challenging for small populations. ConvexGating increasingly outperformed Hypergate with increasing resolution of subpopulations (**Supplementary Fig. 9 C** upper panel left to right and **Supplementary Table 1**). Similar to the small monocyte panel and the large PBMC panel (**Supplementary Fig. 9 A-B**), gating strategies derived by ConvexGating focussed on high precision (**Supplementary Fig. 9 C** mid panel), whereas gating strategies derived by Hypergate prioritized high recall (**Supplementary Fig. 9 C** lower panel). We further compared the performance of ConvexGating to the performance of a linear and an RBF kernel SVM classifier, which directly operated in M-dimensional marker space at the cost of interpretability and direct translatability into a gating strategy (**Supplementary Fig. 10**). ConvexGating outperformed the linear SVM classifier on the DC panel while the linear SVM classifier achieved higher F1 scores for most cell populations on the large PBMC panel and on the cyTOF panel. The non-linear RBF SVM classifier obtained higher F1 scores for nearly all cell populations on the small monocyte panel, the large PBMC panel and on the cyTOF panel. This shows that more flexible machine-learning models find decision boundaries that separate target cells from non-target cells more accurately. However, this comes at the expense of decision boundaries in full marker space that are difficult to interpret and lack direct transferability into a gating strategy which limits application in real-world lab routines.

#### Supplementary Note 3

**Regularized loss.**

Let $C \epsilon R^2$ be the center of the target population $P^t$ and let $H_k$ denote the $kth$ hyperplane. As first regularization component we compute

$$P_1 = \frac{1}{K}\sum_{k=1}^{K} \quad dist(C, H_k)$$

which is the average distance between the hyperplanes and target center $C$.
For all $k = 1, \ldots, K$ $dist$ reads as

$$dist(C, H_k) = \frac{|b_k + w_k^T C|}{||w_k||}$$

where $w_k$ and $b_k$ denote *normal vector* and *bias* of the $kth$ hyperplane. In our second regularization term we consider the location of the target cells relative to the hyperplanes. We exclude those target cells that exhibit the most extreme marker expression profiles in order to reduce the impact of outliers. Let

$$\boldsymbol{q(x) = 1\left[(p_{1,m_1}\ p_{1,m_2})^T \leq x \leq (p_{99,m_1}\ p_{99,m_2})^T\right]}$$

where **1**[·] denotes the indicator function and $p_{u,f}$ the $uth$ percentile of targets $P^t$ along marker $f$. Our second penalty term then reads as

$$P_2(x) = \frac{q(x)}{K}\sum_{k=1}^{K} \quad \left|w_k^T x + b_k\right|.$$

We introduce penalty parameters $\lambda_1 \epsilon R$ and $\lambda_2 \epsilon R$ that control to which extent the regularization components $P_1$ and $P_2$ affect the *loss*. Our *regularized loss* then takes the form

$$\boldsymbol{L_\alpha^{reg}(x) = L_\alpha(x) + \lambda_1 P_1 + \lambda_2 P_2(x)}.$$

For a subset $S \subseteq P^t \cup \quad P^{nt}$ the regularized loss over $S$ reads as

$$L_\alpha^{reg}(S) = \frac{\boldsymbol{1}}{|\boldsymbol{S}|} \ \sum_{\boldsymbol{x \epsilon S}} \quad \boldsymbol{L_\alpha(x) + \lambda_1 P_1} + \frac{\boldsymbol{\lambda_2}}{|\boldsymbol{S}|} \ \sum_{\boldsymbol{x\epsilon S}} \quad \boldsymbol{P_2(x).}$$

### Supplementary figure 1

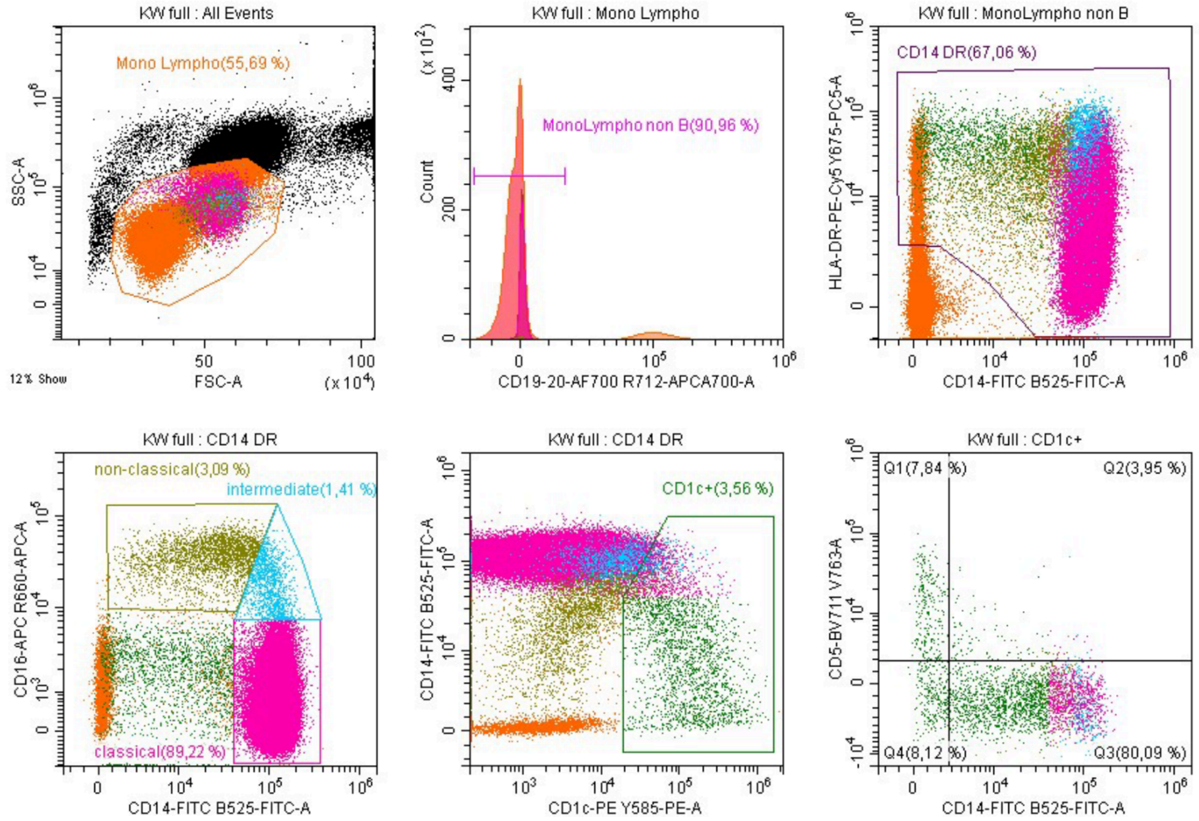


**Overview of the manual gating strategy on the DC panel.** We remove debris (top left panel) and identify B cells (CD19/20+) (top middle panel), monocytes and DC2s (HLA-DR+ or CD14+) (top right panel). We then identify monocyte subtypes (based on CD14 and CD16) (bottom left panel), separate DC2 from monocytes (based CD14 and CD1c) (bottom middle panel), and identify DC2 subtypes (CD1c+, based on CD5 and CD14) (bottom right panel). The gating schema is exemplified for donor ID KW.

### Supplementary figure 2

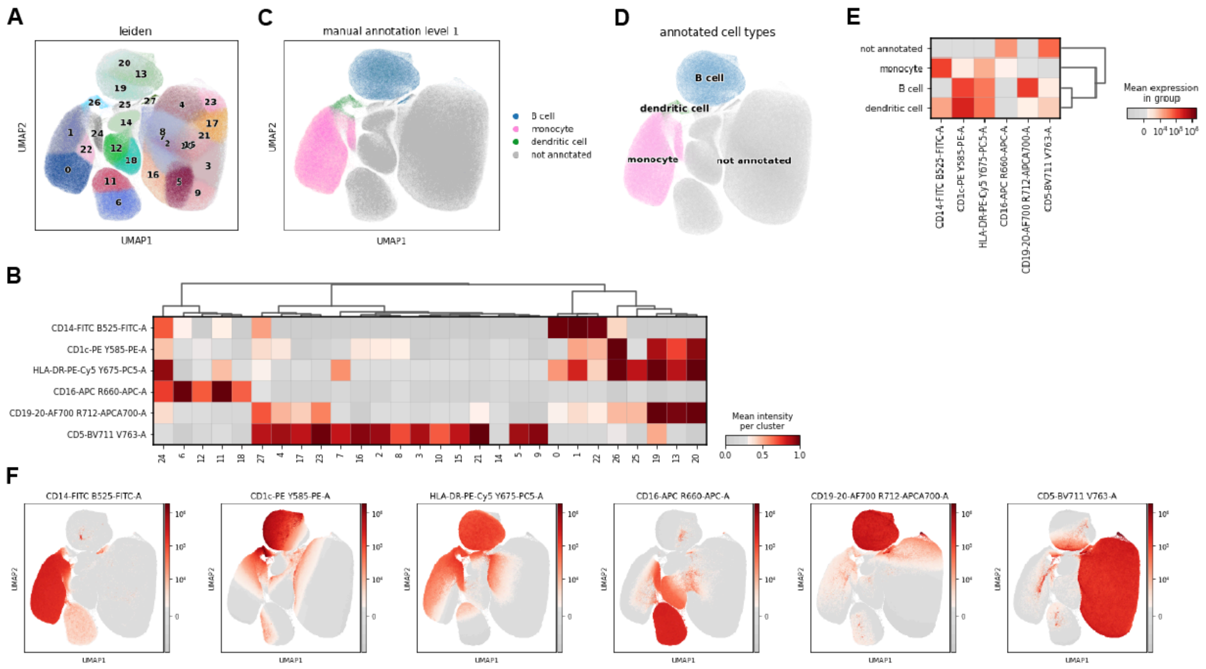

**Leiden clustering and marker based first level annotation. (A)** Leiden clustering of all cells displayed on a UMAP. Clusters are enumerated by size with 0 being the largest and 27 the smallest. **(B)** Mean marker levels per cluster scaled across the entire population. The dendrogram rearranged the clusters by similarity using agglomerative clustering. **(C)** Manual annotation at level 1 displayed on a UMAP. **(D)** First level annotation of clusters to cell types displayed on a UMAP. **(E)** Mean marker levels per cluster across the entire population. **(F)** Marker level distribution for each cell displayed as UMAPs.

### Supplementary figure 3

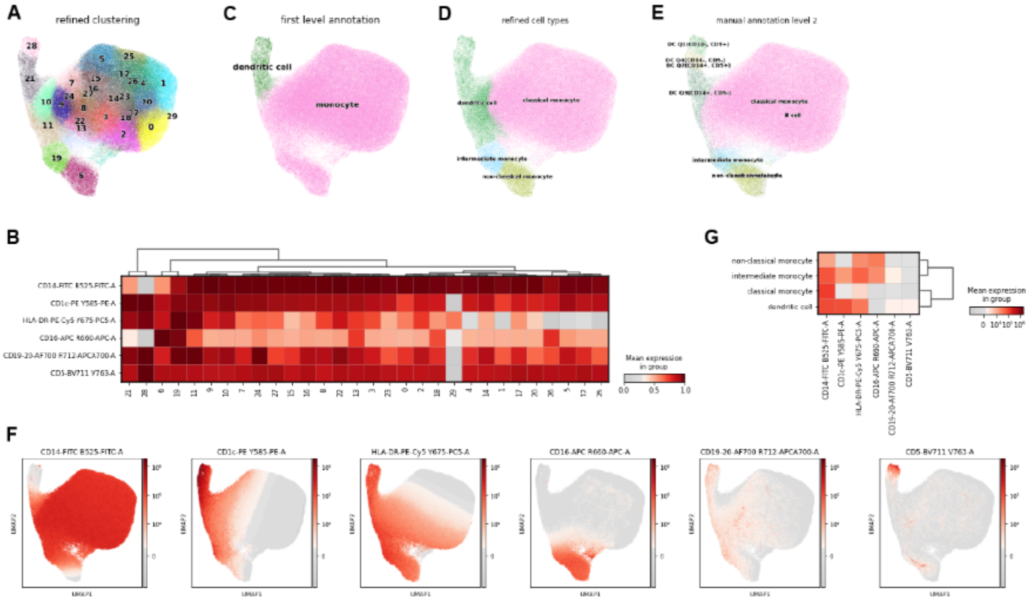


**Leiden clustering and marker based second level annotation for refined monocytes.** **(A)** Leiden clustering for cells annotated as monocytes and DC2s displayed on a UMAP. Clusters are enumerated by size with 0 being the largest and 29 the smallest. **(B)** Mean marker levels per cluster scaled across the monocyte/DC2 subpopulation. The dendrogram rearranged the clusters by similarity using agglomerative clustering. **(C)** First level annotation of clusters to cell types displayed on a UMAP. **(D)** Refined annotation of clusters to monocyte subtypes and DC2s displayed on a UMAP. **(E)** Manual annotation at level 2 displayed on a UMAP. **(F)** Marker level distribution for each cell displayed as UMAPs. **(G)** Mean marker levels per cluster across the monocyte/DC2 subpopulation.

### Supplementary figure 4

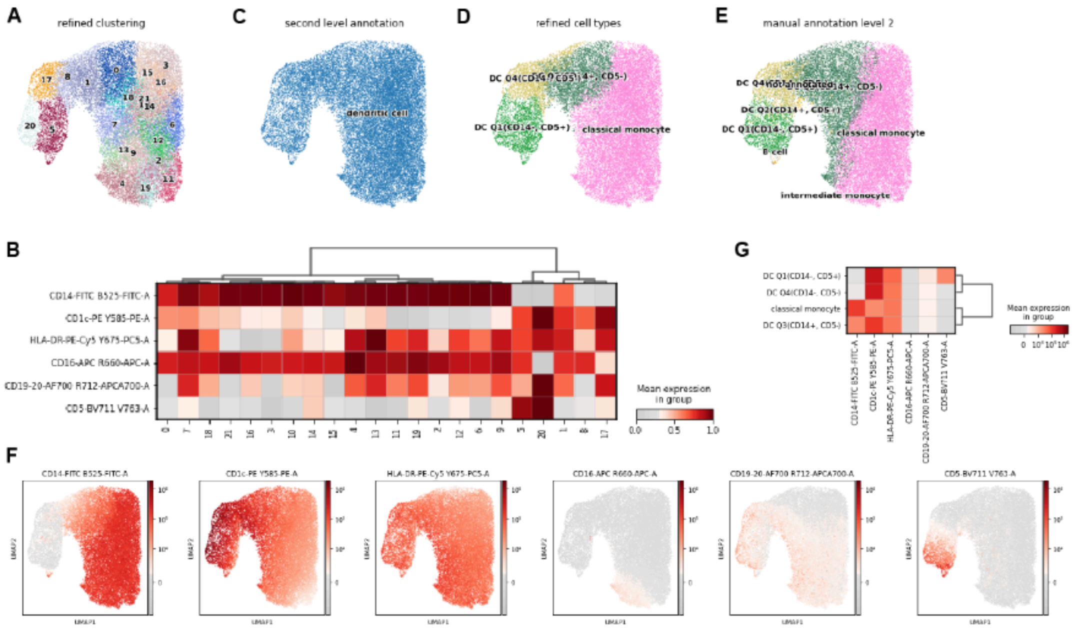


**Leiden clustering and marker based second level annotation for refined DC2s. (A)** Leiden clustering for cells annotated as DC2s displayed on a UMAP. Clusters are enumerated by size with 0 being the largest and 20 the smallest. **(B)** Mean marker levels per cluster scaled across the monocyte/DC2 subpopulation. The dendrogram rearranged the clusters by similarity using agglomerative clustering. **(C)** Second level annotation of clusters to cell types displayed on a UMAP. **(D)** Refined annotation of clusters to monocyte subtypes and DC2s displayed on a UMAP. **(E)** Manual annotation at level 2 displayed on a UMAP. **(F)** Marker level distribution for each cell displayed as UMAPs. **(G)** Mean marker levels per cluster across the monocyte/DC2 subpopulation.

### Supplementary figure 5

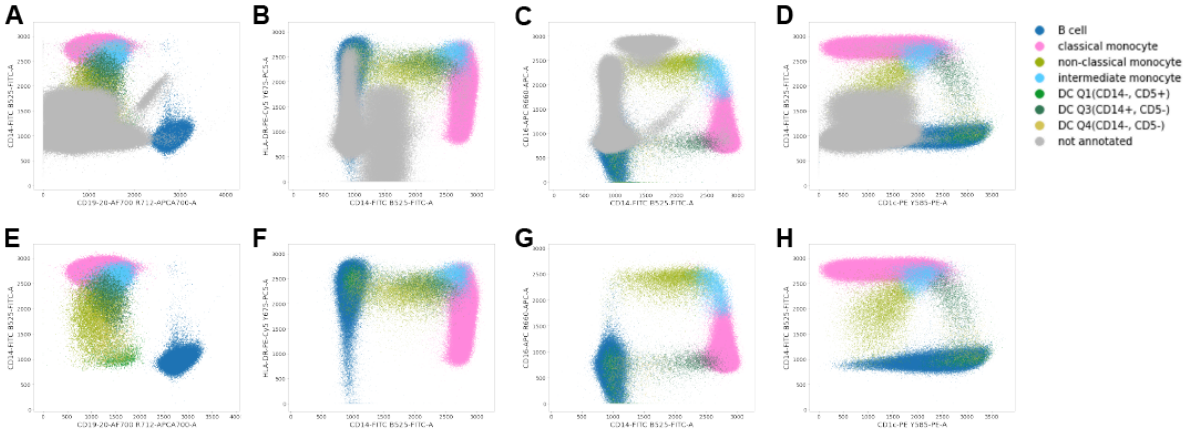


**2D scatter plots of the Leiden clustering based cell type annotation using the DC panel for whole blood staining of three donors. (A-D)** All cells colored by annotation level 2 including not annotated cells. (**E-H**) All annotated cells colored by annotation level 2 excluding not annotated cells.

### Supplementary figure 6

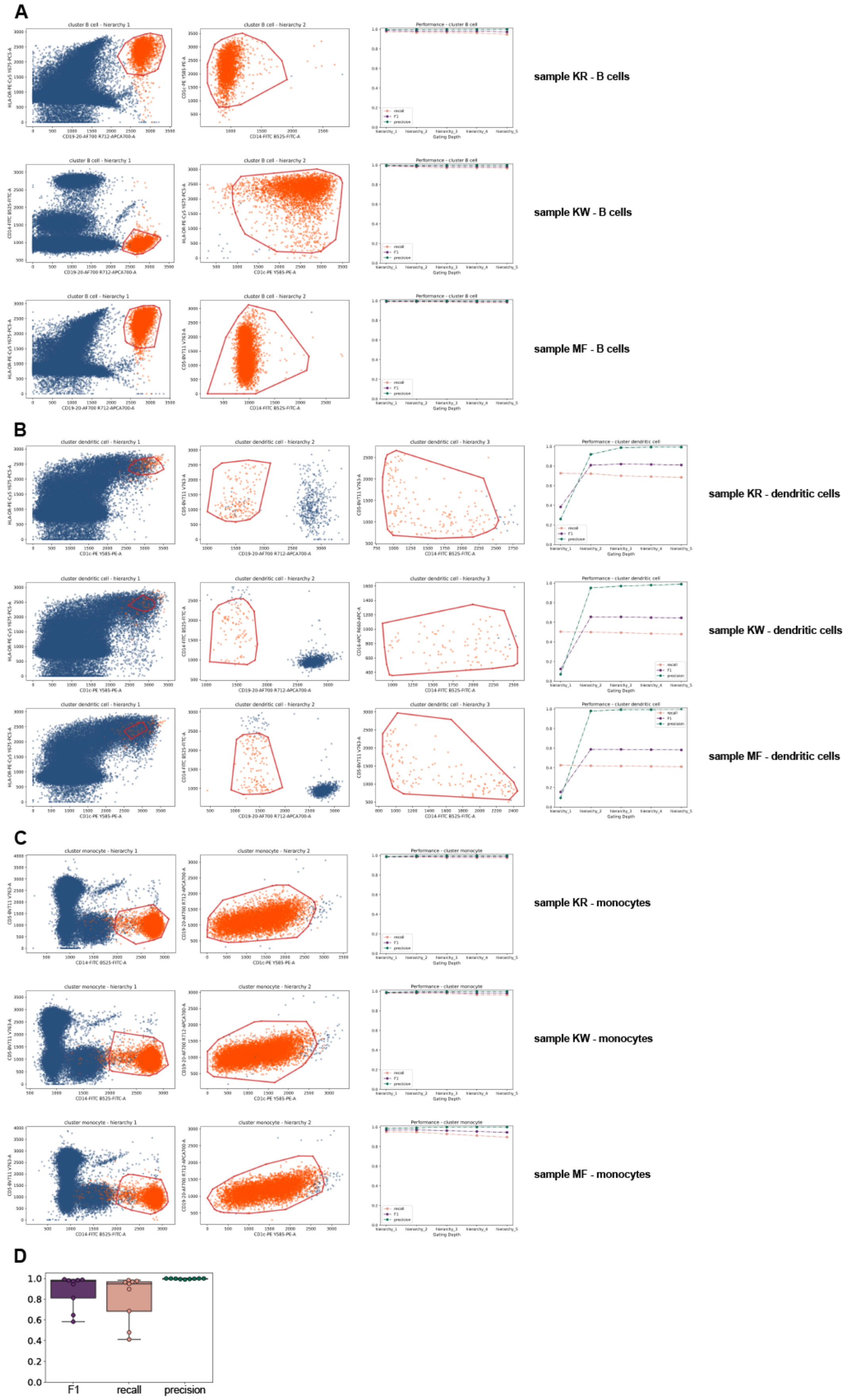

**E**

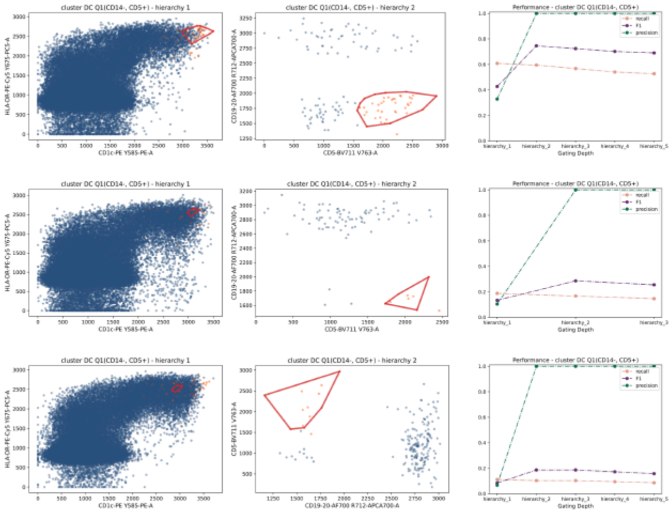

sample KR - dendritic cell Q1 (CD14-, CD5+)

sample KW - dendritic cell Q1 (CD14-, CD5+)

sample MF - dendritic cell Q1 (CD14-, CD5+)

**F**

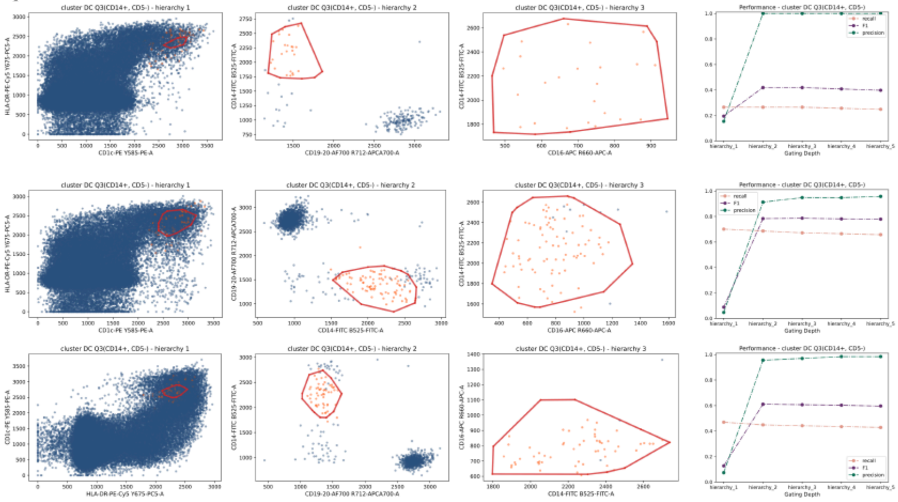

sample KR - dendritic cells Q3 (CD14+, CD5-)

sample KW - dendritic cells Q3 (CD14+, CD5-)

sample MF - dendritic cells Q3 (CD14+, CD5-)

**G**

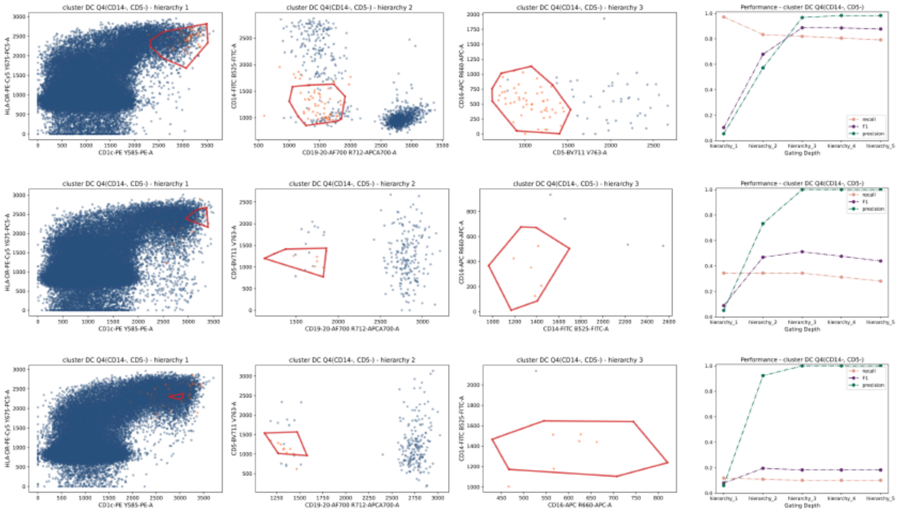

sample KR - dendritic cells Q4 (CD14-, CD5-)

sample KW - dendritic cells Q4 (CD14-, CD5-)

sample MF - dendritic cells Q4 (CD14-, CD5-)

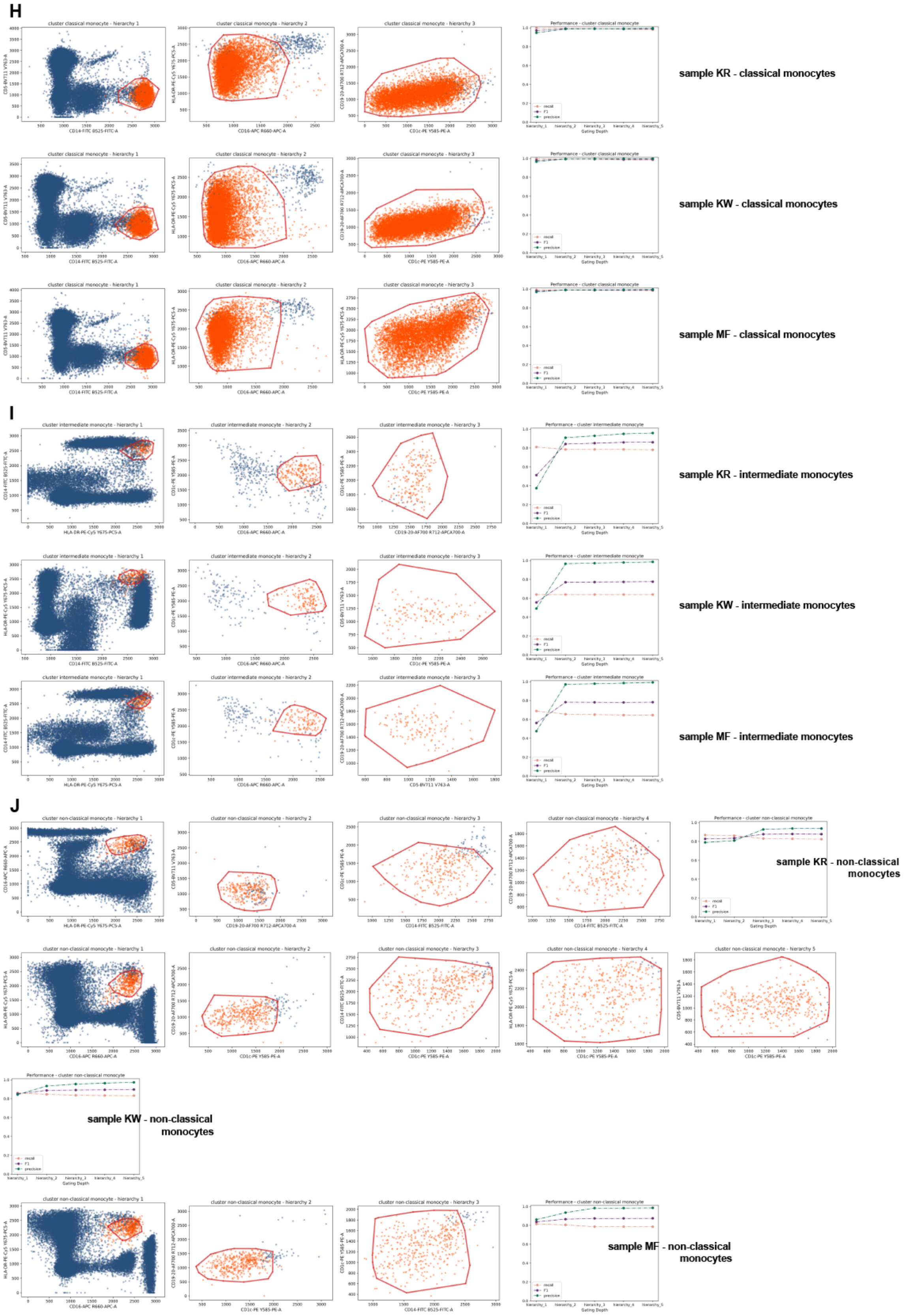


**Inferred gating strategies for the DC panel. (A-C)** Gating strategies inferred by ConvexGating for cell type annotation level 1. **(D)** Boxplot showing precision, recall and F1 for all samples in cell type annotation level 1. **(E-J)** Gating strategies inferred by ConvexGating for cell type annotation level 2.

### Supplementary figure 7

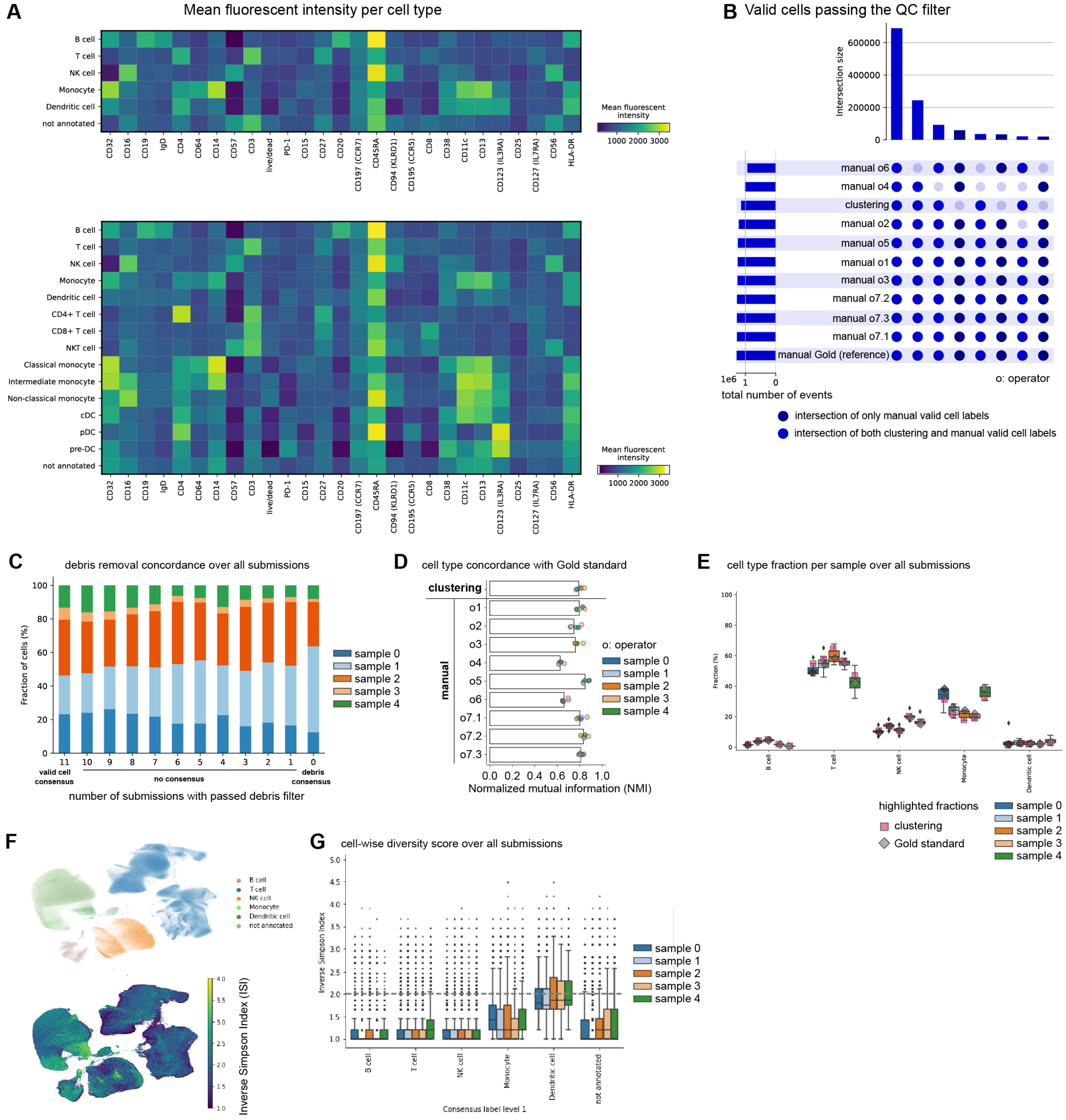

H

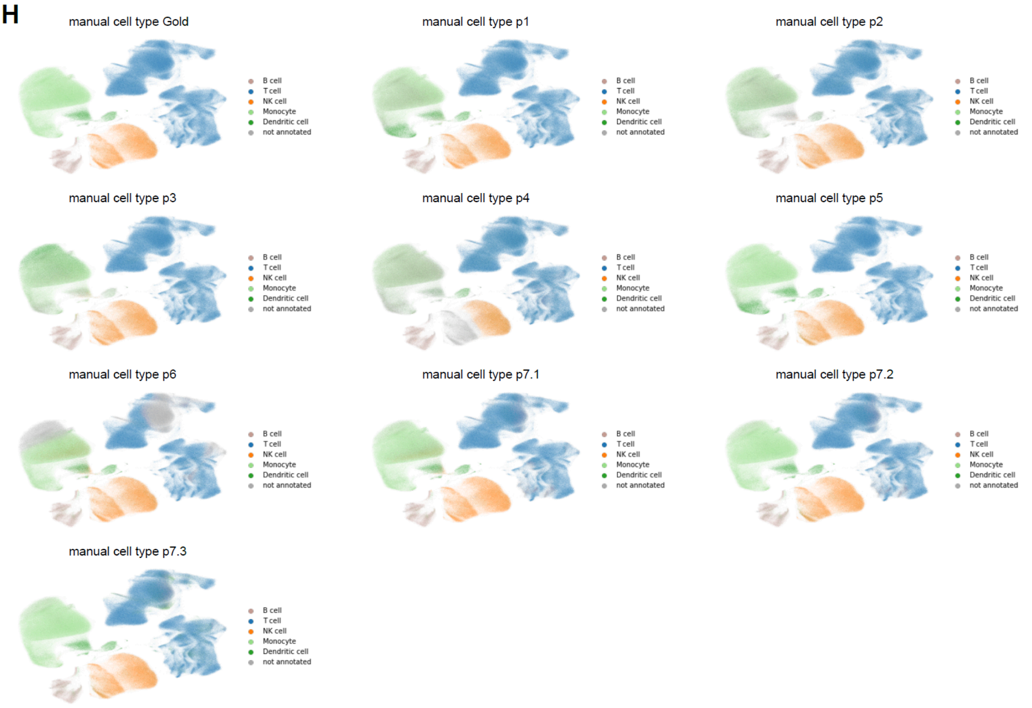
manual cell type Gold
manual cell type p1
manual cell type p2
manual cell type p3
manual cell type p4
manual cell type p5
manual cell type p6
manual cell type p7.1
manual cell type p7.2
manual cell type p7.3
B cell
T cell
NK cell
Monocyte
Dendritic cell
not annotated

I

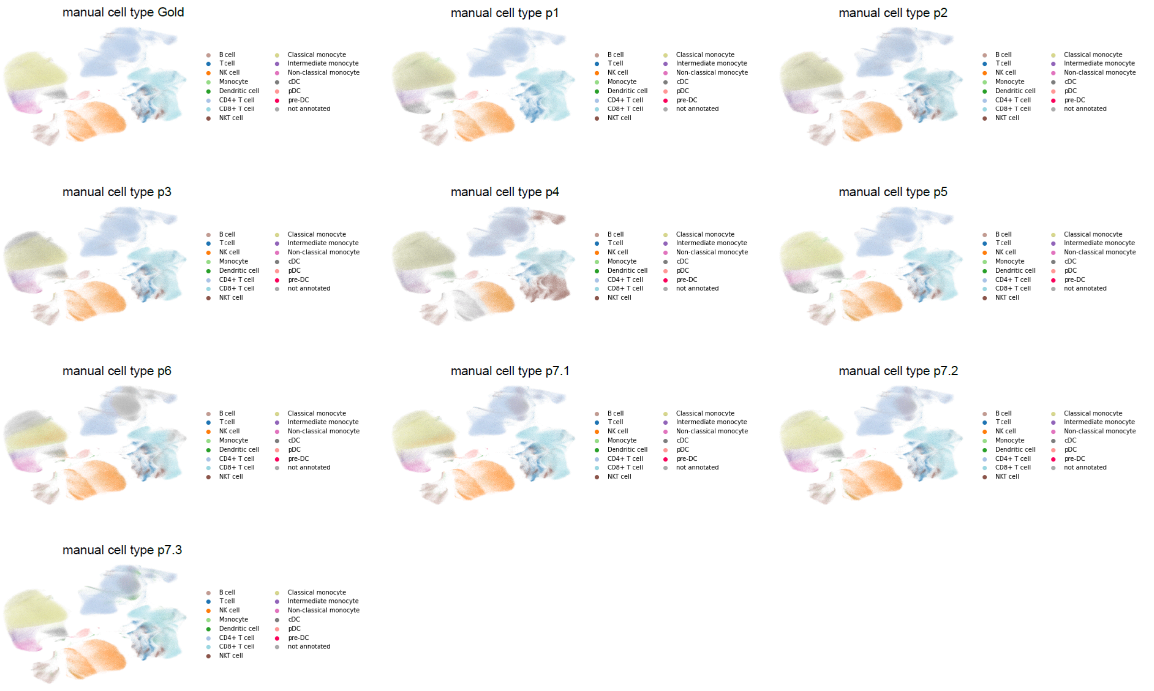
manual cell type Gold
manual cell type p1
manual cell type p2
manual cell type p3
manual cell type p4
manual cell type p5
manual cell type p6
manual cell type p7.1
manual cell type p7.2
manual cell type p7.3
B cell
T cell
NK cell
Monocyte
Dendritic cell
CD4+ T cell
CD8+ T cell
NKT cell
Classical monocyte
Intermediate monocyte
Non-classical monocyte
cDC
pDC
pre-DC
not annotated

J

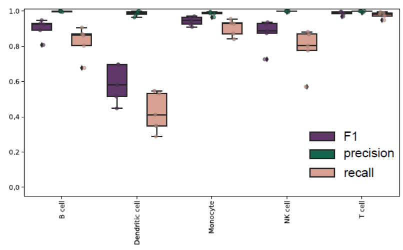
1.0
0.8
0.6
0.4
0.2
0.0
B cell
Dendritic cell
Monocyte
NK cell
T cell
F1
precision
recall

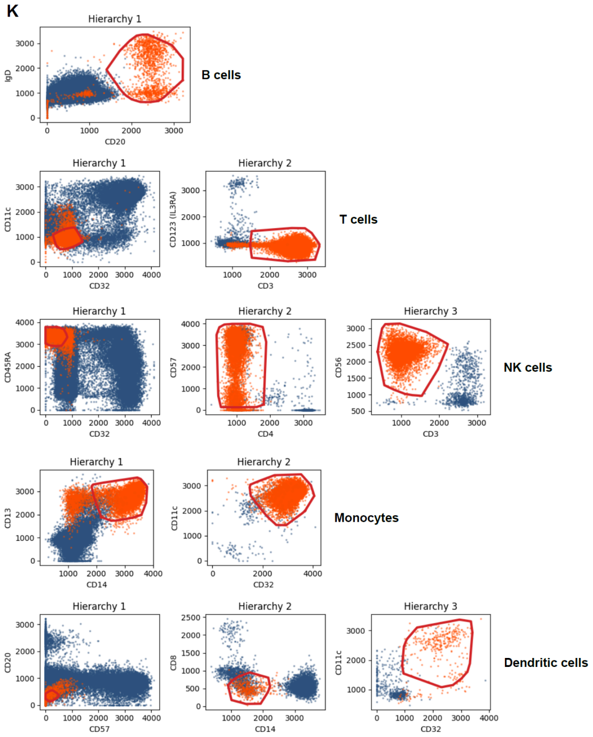
K
B cells
T cells
NK cells
Monocytes
Dendritic cells

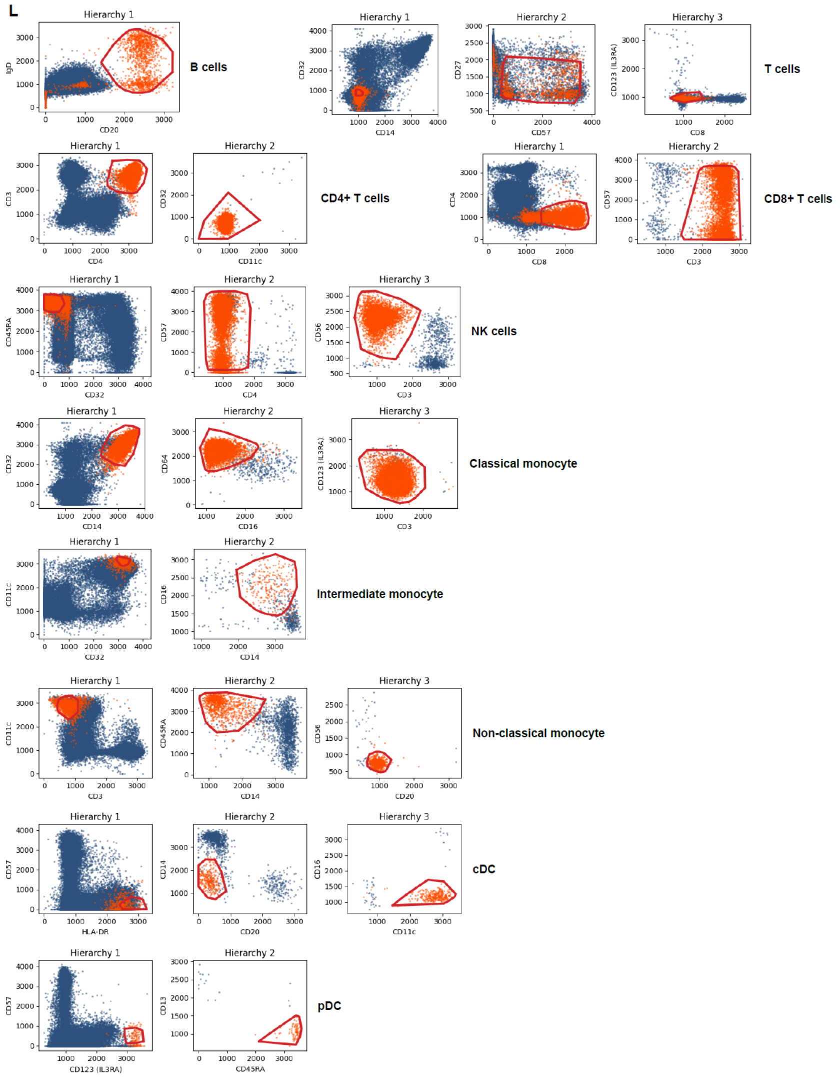
L
B cells
T cells
CD4+ T cells
CD8+ T cells
NK cells
Classical monocyte
Intermediate monocyte
Non-classical monocyte
cDC
pDC

**Detailed variability of human expert labeled data. (A)** Mean fluorescent intensity of all 27 markers per consensus cell type (upper panel: coarse cell type level 1, lower panel: finer cell type level 2). **(B)** Upsetplot of the overlap of cells passing the QC filter ("valid cells") across all submissions. **(C)** Fraction of valid cells per sample aggregated over all submissions. Cells consistently passing QC have highest scores. We do not observe a sample bias in the QC step. **(D)** Barplot of overlap of cell type labels for cell type level 1 compared to Gold standard measured by normalized mutual information (NMI). Dots colored by sample indicate NMI per sample. **(E)** Boxplot of cell type composition (cell type level 1) over all submissions(n=11, boxes represent 25th percentile, median and 75th percentile, respectively, whiskers show 1.5 * interquartile range and black diamonds represent outliers), colored by sample. Gray squares indicate cell type annotation by clustering, gray diamonds indicate Gold standard annotation. **(F)** UMAPs displaying the consensus cell type labels (cell type level 1) over all submissions and Gold standard (upper panel) and the number of effective labels per cell over all human expert labels measured by Inverse Simpson Index (ISI) (lower panel). **(G)** Boxplot of number of effective labels per cell over all submissions(n=11, boxes represent 25th percentile, median and 75th percentile, respectively, whiskers show 1.5 * interquartile range and black diamonds represent outliers), colored by sample, grouped by the consensus label over all submissions and the Gold standard. Grey dashed horizontal line indicates the effective number of 2 cell type labels. **(H)** UMAPs of all expert label submissions, the Gold standard and clustering for cell type level 1. **(I)** UMAPs of all expert label submissions, the Gold standard and clustering for cell type level 2. **(J)** Boxplots summarizing precision, recall and F1 scores per sample obtained by running ConvexGating on all samples for cell populations annotated by clustering on cell type level 1. **(K)** Gating strategies inferred by ConvexGating for cell type level 1. **(L)** Gating strategies inferred by ConvexGating for cell type level 2.

### Supplementary figure 8

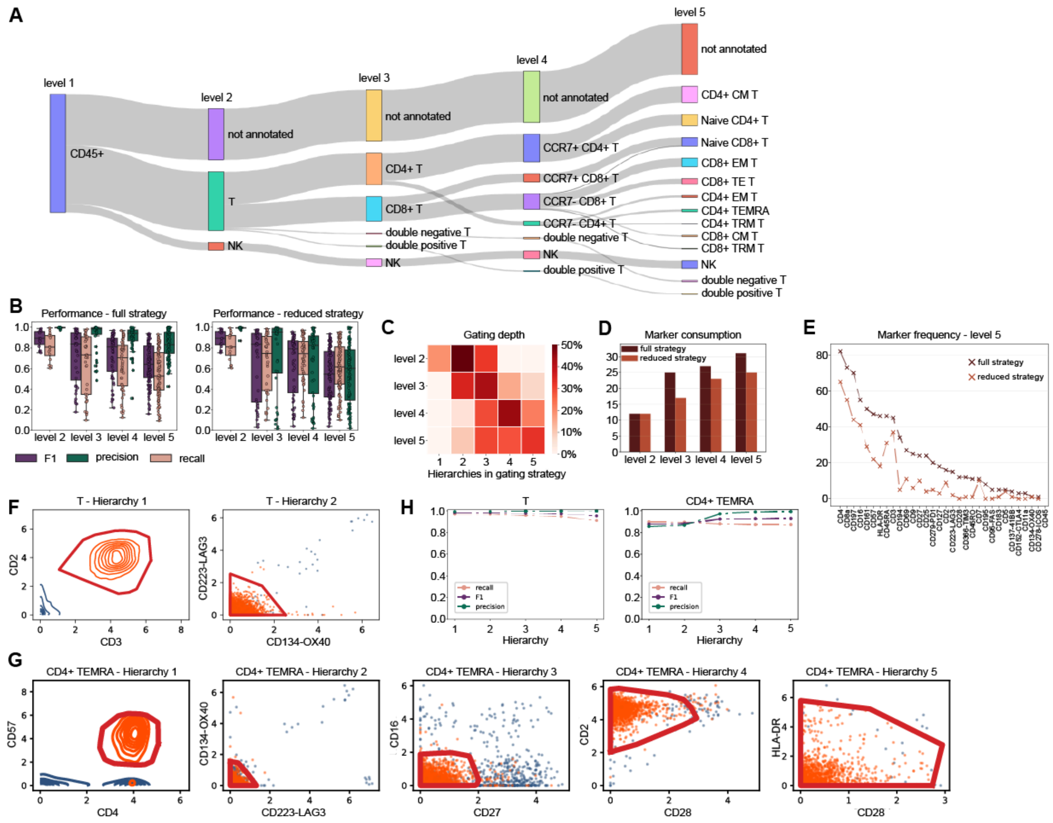

**ConvexGating finds gating strategies for cell populations (four annotation levels, eight samples) in healthy human bone marrow**[33]. **(A)** Sankey plot visualizing T cell subtyping with increasing annotation level. **(B)** Performance overview (F1, recall, precision) per cell type annotation level for the full gating strategies (left panel) and for the gating strategies consisting of the initial two gating hierarchies (reduced strategies, right panel) (boxes represent 25th percentile, median and 75th percentile, respectively, whiskers show 1.5 * interquartile range and black diamonds represent outliers). Dots indicate the performance score for a specific cell type and sample. **(C)** Distribution of hierarchy depth in gating strategies inferred by ConvexGating per cell type annotation level. **(D)** Marker usage for gating all subpopulations in all samples per cell type annotation level. The left bar (dark red) shows the marker usage for the full gating strategies while the right bar (light red) shows the marker usage for the reduced strategies. **(E)** Frequency of marker choice in gating strategies for cell type annotation level 5. Dark red crosses refer to the full gating strategies while light red crosses refer to the reduced gating strategies. **(F)** Visualization of gating strategy for T cells (annotation level 2, sample A) retrieved by ConvexGating. **(G)** Visualization of the gating strategy for CD4+ TEMRA cells (annotation level 5, sample A). Target cells (orange) and non-target cells (blue) are represented via scatter plots and/or contour plots per target and non-target population. **(H)** Overview of ConvexGating performance (F1 score, recall, precision) for T cell cluster (annotation level 2, sample A) and CD4+ TEMRA cell cluster (annotation level 5, sample A).

### Supplementary figure 9

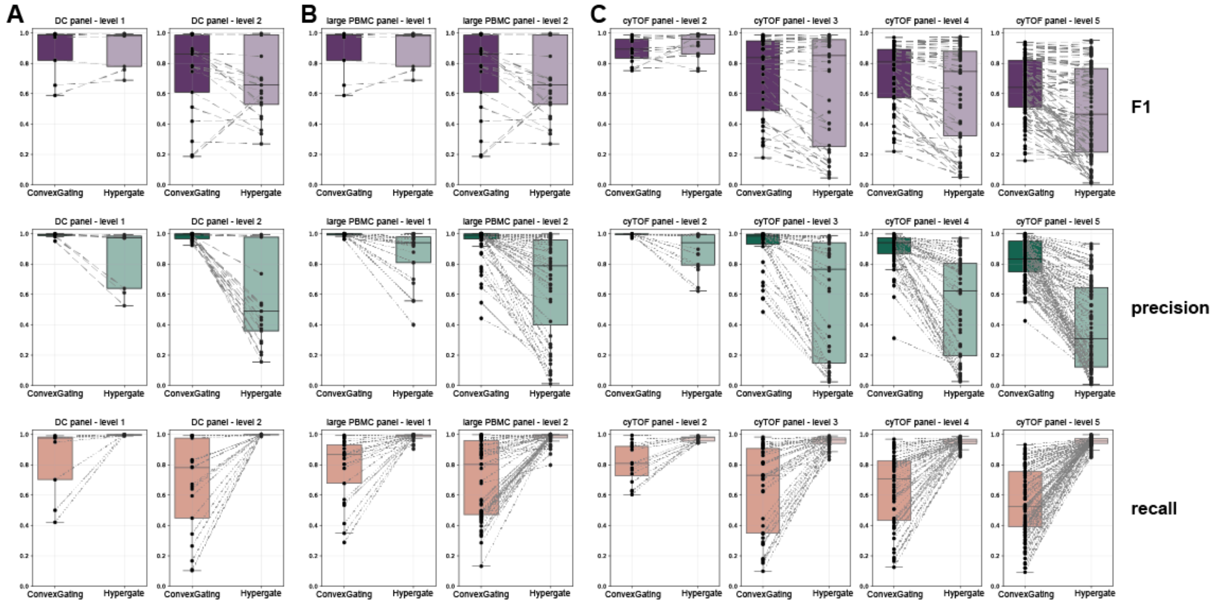


**Computational benchmark analysis of gating strategies derived by ConvexGating versus gating strategies derived by Hypergate**. F1 (upper panel), precision (mid panel) and recall (lower panel) boxplot results for gating strategies learned by ConvexGating (darker shade) and Hypergate (lighter shade). Boxes represent 25th percentile, median and 75th percentile, respectively, whiskers show 1.5 * interquartile range and black diamonds represent outliers. Each dot represents the gating performance for a specific cell subpopulation. The dotted gray lines connect the corresponding subpopulations in ConvexGating and Hypergate boxplots. The multimodal benchmark analysis includes a DC panel with two cell type annotation levels **(A)**, a large PBMC panel with two cell type annotation levels **(B)** and a cyTOF panel with four annotation levels **(C)**.

### Supplementary figure 10

Performance of linear SVM

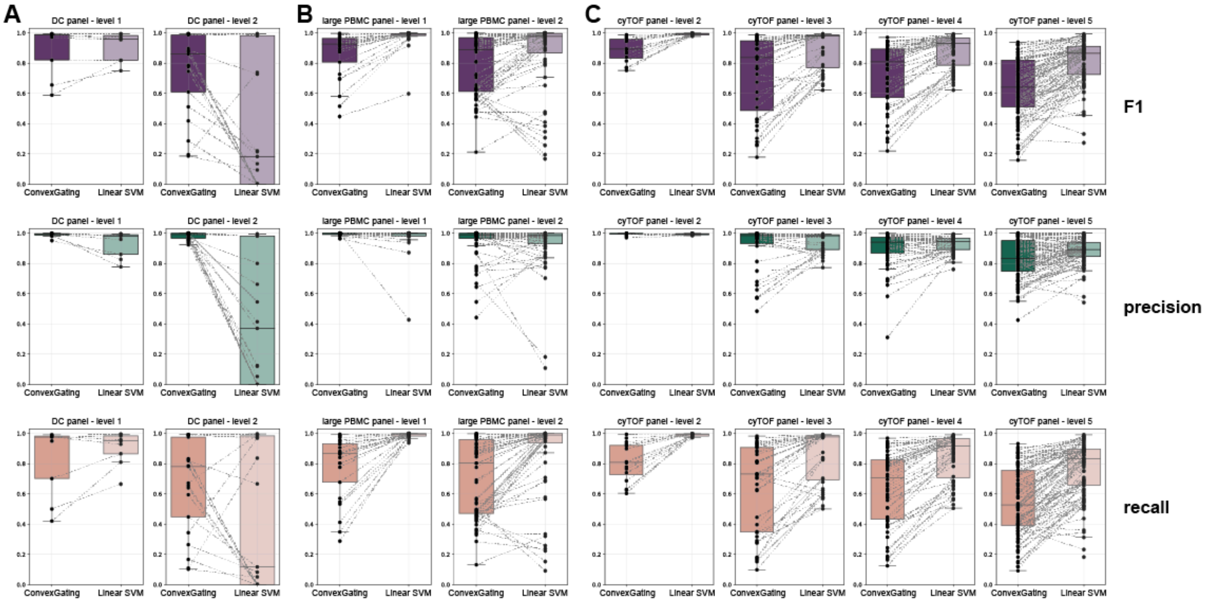


Performance of RBF kernel SVM

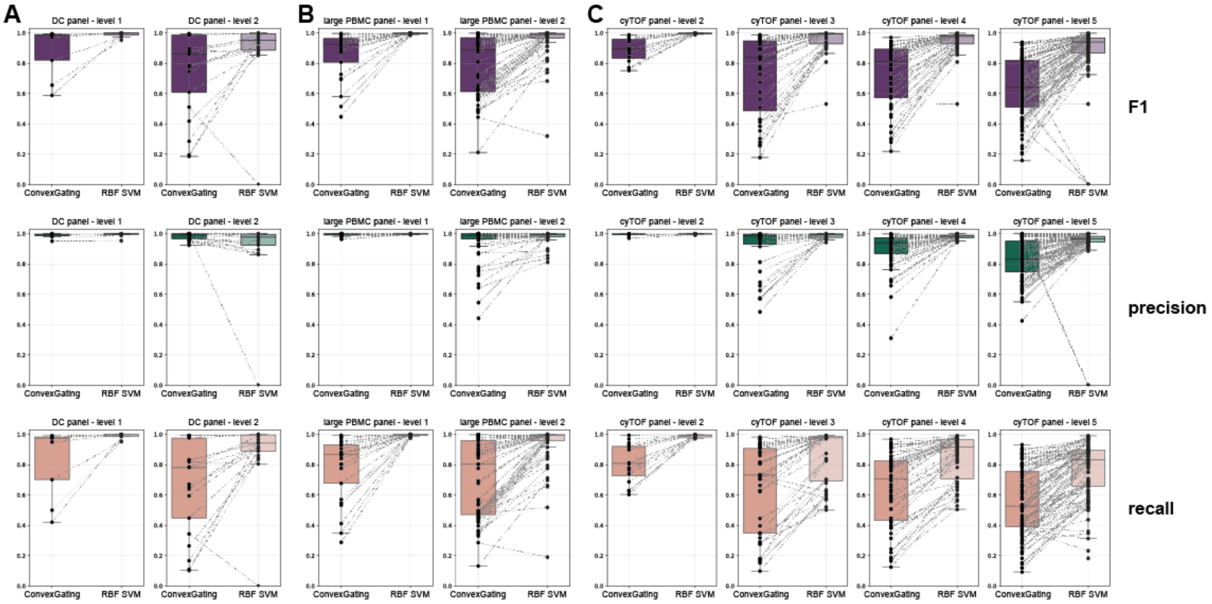


**Computational benchmark analysis of gating strategies derived by ConvexGating versus performance of linear SVM (upper panel) and RBF kernel SVM (lower panel).** The multimodal benchmark analysis includes a DC panel with two cell type annotation levels **(A)**, a large PBMC panel with two cell type annotation levels **(B)** and a cyTOF panel with four annotation levels **(C)**. Boxes represent 25th percentile, median and 75th percentile, respectively, whiskers show 1.5 * interquartile range and black diamonds represent outliers. Each dot represents the gating performance for a specific cell subpopulation. The dotted gray lines connect the corresponding subpopulations in ConvexGating and Hypergate boxplots.

### Supplementary figure 11

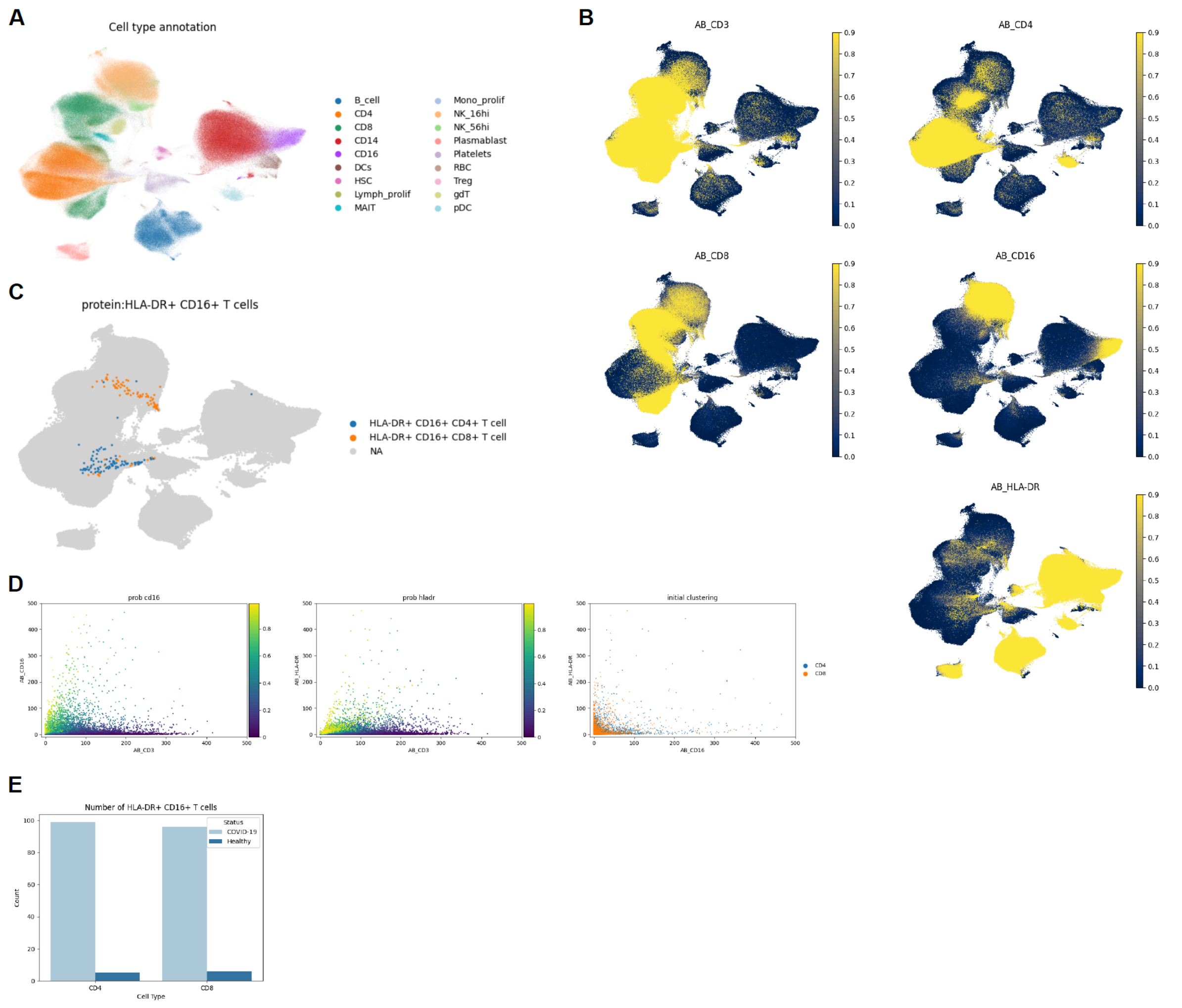


**Identification of CD16+ T cells in COVID-19 CITEseq data[34]. (A-C)** UMAPs showing initial cell type annotation[34] **(A)**, foreground probability of the markers CD3, CD4, CD8, CD16, and HLA-DR **(B)**, and CD16+ T cell annotation **(C)**. **(D)** Scatter plot of drop-in corrected anti-CD16, anti-HLA-DR and anti-CD3 abundance in the T cell compartment. Dots in the left panels are colored by foreground probability. Dots in the right panel are colored by cell type. **(E)** Absolute number of HLA-DR CD16+ T cells per disease diagnosis over all samples.
